# SRSF3 is oncogenic in breast but tumor-suppressive in liver by differential regulation of gene expression

**DOI:** 10.1101/2025.03.14.643315

**Authors:** Lulu Yu, Masahiko Ajiro, Alexei Lobanov, Maggie Cam, Vladimir Majerciak, Baktiar Karim, Deanna Gotte, Chu-Xia Deng, Douglas R. Lowy, Nicholas J.G Webster, Zhi-Ming Zheng

**Author notes:** Co-first authors (Lulu Yu, Masahiko Ajiro). Division of Cancer RNA Research, National Cancer Center Research Institute, Tokyo 104-0045, Japan. Correspondence: Zhi-Ming Zheng.

## Abstract

SRSF3 (SRp20) is an essential splicing factor. We discovered Srsf3 plays an oncogenic role in breast cancer and Srsf3 knockout (KO) in mammary glands delays the development of breast cancer in an Erbb2 mouse model. In contrast, Srsf3 is tumor-suppressive in mouse liver tissues. Srsf3 KO in hepatocytes enhances DEN-induced liver cancer and disrupts the sex disparity in DEN-induced liver cancer. Comparing to Srsf3 WT liver cancer, Srsf3 KO significantly increases Sox4, E2f1, Trpv4, Trim6, and Myc expression, but does not so in Erbb2 breast cancer. Srsf3 KO inhibits expression of Mfsd4a and Eif4a2 in breast cancer but enhances Mfsd4a and Eif4a2 expression in liver cancer. Moreover, Srsf3 KO suppresses the expression of ERα and Foxa genes to reduce Lifr and Egfr but induce Myc expression and promote liver cancer in female mice. Together, our data highlight a new functional paradigm of SRSF3 at its physiological level in tissue context-dependent gene regulation.

## Introduction

Serine and arginine rich splicing factor 3 (SRSF3), also called SRp20 or SFRS3, is the smallest member of key splicing factors, serine and arginine rich proteins (SR proteins) involved in regulating pre-mRNA splicing (1–4). SRSF3 promotes RNA splicing when binding to an exonic cis-element (5–9), but inhibits RNA splicing when binding to an intronic element (10). Besides mRNA splicing, SRSF3 as a cytoplasm-nuclear shuttling protein plays critical roles in many cellular processes, including RNA polyadenylation (11,12), RNA export (13,14), miRNA biogenesis (7,15,16) and protein translation (17,18). SRSF3 was found indispensable for embryo development (19). Moreover, we recently discovered that SRSF3 is an anti-apoptotic and senescent factor essential for cell proliferation by promoting the G2/M transition and is oncogenic as a proto-oncogene when overexpressed (20,21) by regulating genome-wide expressions of numerous genes at transcriptional and post-transcriptional levels (7,22). A variety of human cancer tissues exhibit highly increased expression of SRSF3 (4,20,23,24). In contrast, ubiquitous knockout of Srsf3 results in embryonic lethality due to a failure in blastocyst formation (19).

Breast cancer is the most frequently diagnosed cancer and a leading cause of cancer death in women worldwide (25). Breast cancer tissues show increased SRSF3 expression (20) (Figure S1), and the increased expression of SRSF3 was found to correlate with breast cancer progression and 5-year overall survival (20). SRSF3 was found to be essential for triple-negative breast cancer progression by interacting with TDP43 and regulation of alternative RNA splicing of CD44 variants (26,27). Nevertheless, the function of SRSF3 in other types of breast cancer remains to be explored.

Liver cancer is the sixth most commonly diagnosed cancer and ranks third in cancer mortality worldwide (25). SRSF3 expression is high in most liver cancer cases (20) (Figure S1). SRSF3 is essential for the differentiation of hepatocytes and their metabolic function (28). SRSF3 reduction contributes to progressive liver disease (29), and mice with SRSF3 KO developed liver cancer with aging (30), suggesting a possible tumor suppressive effect of SRSF3. However, the molecular mechanisms of how Srsf3 function at physiological level and in the development of liver cancer remains to be determined.

Both human SRSF3 gene in size of 8422 bp on chromosome 6 and mouse Srsf3 gene in size of 9423 bp on chromosome 17 contain seven exons and six introns and encode 164-aa SRSF3 protein with 100% homology between the mouse and human. Their transcription start sites and polyadenylation cleavage sites were recently remapped and updated (31). The major SRSF3 mRNA isoform encoding the full-length protein is 1411-nt long generated by alternative RNA splicing and alternative polyadenylation with exclusion of exon 4 (a poison exon) (31). Similarly, major mouse Srsf3 mRNA isoform is only 1295-nt long with a shorter 3’ UTR over the human SRSF3 3’ UTR. The sizes of both human and mouse SRSF3 mRNAs aremuch shorter than the NCBI annotated SRSF3 transcripts (31). By using Cre-LoxP technology and MMTV promoter-driving Cre expression in mammary tissues and Albumin (Alb)-promoter-driving Cre expression in liver tissues in the current study, we conditionally knocked out Srsf3 expression separately from mouse breast and liver tissues. Srsf3 knockout (KO) on breast carcinogenesis was evaluated by further crossing the Srsf3 knockout mouse with the mouse carrying a mutant c-neu (Erbb2 or HER2)(32). Srsf3 KO on liver carcinogenesis was evaluated by intraperitoneal injection of 15 days-old mice with diethylnitrosamine (DEN), a chemical liver carcinogen (33). We demonstrated that Srsf3 is oncogenic in breast cancer formation, but tumor-suppressive in liver cancer formation. Using genome-wide analyses, we discovered a distinguished set of gene expression profiles from the breast cancer to liver cancer.

## Materials and Methods

### Generation of transgenic mice

Srsf3^tm1Pjln^ mice (Srsf3^flox/flox^ BALB/c) were provided from Hassan Jumaa (19). Albumin promoter-regulated Cre (Alb-Cre) expressing strain (C57BL/6) and MMTV promoter-regulated Cre (MMTV-Cre) expressing strain (FVB) were provided from R.-H.W. and C.-X. D. MMTV promoter-regulated rat mutant c-neu strain, FVB-Tg (MMTV-Erbb2) NK1Mul/J, was purchased from the Jackson Laboratory (Bar Harbor, ME). NCI-Frederick is accredited by AAALAC International and follows the Public Health Service Policy for the Care and Use of Laboratory Animals. Animal care was provided in accordance with the procedures outlined in the ‘Guide for Care and Use of Laboratory Animals (National Research Council; 1996; National Academy Press; Washington, D.C.).

### Induction and monitoring tumor formation

Liver cancer formation was induced by i.p. injection of diethylnitrosamine (DEN) (25 mg/kg of body weight) at 15 days of age (33). Breast cancer formation was induced by c-neu expression from MMTV promoter. Weekly survey for liver and breast tumor formation were conducted by palpation by facility veterinarian in a blind manner for individual genotypes up to 78 weeks. Animals presenting with tumors exceeding 20 mm in dimension, necrotic tumors, cutaneous ulceration, or multiple tumors collectively weighing more than 10% of the animal’s body weight were euthanized via CO2 asphyxiation. This procedure was conducted at humane endpoints in accordance with institutional Animal Care and Use Committee (ACUC) guidelines for tumor collection and biochemical analysis. Breast tissues and liver tissues at age of 6 months and terminal life stage were collected for pathology analysis.

### Study approval

Our animal study was approved by the Institutional Animal Care and Use Committee of NCI-Frederick (protocol numbers 14-101, 17-101, 19-101, 22-101) and of NIH (HAMB-001).

### Total RNA sequencing and data processing

Total RNA was extracted from the collected Srsf3 WT breast/liver tumors, Srsf3 KO breast tumors and Srsf3 KO liver tumors using TriPure reagent (Roche, # 11667165001). For liver tumor tissues RNA-seq, paired-end 125-bp read length sequencing with a depth of 100 million reads per sample was performed on the Illumina platform according to the manufacturer’s instructions. For breast tumor tissues RNA-seq, the sequencing libraries were constructed following Illumina Stranded Total RNA protocol (Illumina, RS-122-2201). The length of the paired-end read was 150-bp with a depth of 100 million reads per sample. The obtained sequence reads in fastq format were mapped to the mouse reference genome (mm10), and Limma-voom was used for quantile normalization and calculation of differentially expressed genes. Genes with a p-value ≤0.05, and adjustment of p-values by Benjamini Hochberg FDR (FDR ≤0.05), were considered statistically significant differentially expressed. rMATs (34) software was used for differential alternative RNA splicing analysis. Instant Clue software (version 0.12.1) and ClustVis online heatmap software (https://biit.cs.ut.ee/clustvis/) were used to generate the volcano plots and heatmaps, respectively.

### NanoString nCounter gene expression validation

Four samples from four groups in each tumor model, including Srsf3 WT, Srsf3 homogenous knockout mice at 6^th^ month and tumor stage in both breast and liver group, were submitted for NanoString nCounter gene expression analysis. Top upregulated or downregulated genes as well as genes not in top list, but have important functions identified by RNA-seq analysis in breast and liver tumors were chosen for validation (Table S3). Probe sets for breast tissues and liver tissues were custom designed and synthesized by NanoString company. Raw counts were extracted and normalized to three chosen housekeeping genes using nSolver 3.0 digital analyzer software.

### Cell lines and siRNAs

Mouse liver cancer cell Hepa1-6 and human liver cancer cell Huh7 was obtained from T. Jake Liang’s lab (NIDDK/NIH). Mouse breast cancer cell NF639 expressing c-neu oncogene, human breast cancer cell SKBR3 (HER2+) and MCF7 (ER+) was purchased from ATCC. SKBR3 cell was maintained in McCoy’s 5A Medium (Thermo Fisher Scientific) with 10% fetal bovine serum (FBS, Cytiva). All the other cell lines were maintained in Dulbecco’s modified Eagle’s medium (DMEM) (Thermo Fisher Scientific) with 10% FBS at 37°C under a 5% CO_2_ atmosphere.

For siRNA transfection, all the cells at 24 h of cell passage were transfected with 40 nM of siRNA, and total protein and RNA was extracted from the harvested cells 48 h after the transfection. LipoJet In Vitro Transfection Kit (Ver. II) (SignaGen Laboratories, # SL100468) was used for Hepa1-6 cell, Huh7 cell and MCF7 cell transfection. Lipofectamin RNAiMAX Transfection Reagent (Thermo Fisher Scientific, # 13778075) was used for NF639 cell transfection. SKBR3 transfection kit (Altogen Biosystems, #6905) were used for SKBR3 cell transfection. siRNAs from Dharmacon used in this study: mouse Srsf3 (# M-059214-01) and human SRSF3 (# M-030081-00). Non-targeting control siRNA (# D-001210-01) served as a negative control.

### RT-PCR and RT-qPCR

For reverse transcription-coupled polymerase chain reaction (RT-PCR), 1 µg of DNase-treated total RNA was reverse transcribed with MuLV RTase and random hexamer (Thermo Fisher Scientific), and PCR-amplified with AmpliTaq (Thermo Fisher Scientific). Primer sets used for RT-PCR were listed in Table S6. Quantitative RT-PCR (RT-qPCR) was conducted by TaqMan real-time PCR system (Thermo Fisher Scientific) for Erα (# Mm00433149_m1), Sox4 (# Mm00486320_s1), E2f1(# Mm00432939_m1),Trpv4 (# Mm00499025_m1), Trim6 (# Mm07305537_m1), Foxa1 (# Mm00484713_m1), Foxa2 (# Mm00839704_mH), Foxa3 (# Mm00484714_m1), Lifr (# Mm00442942_m1), Egfr(# Mm01187858_m1), Myc (# Mm00487804_m1), Htatip2 (#Mm00457476_m1) and Gapdh (# Mm99999915_g1).

### Western blot and immunohistochemistry (IHC)

For Western blot, SDS-PAGE was performed in 4-12% Bis-Tris NuPAGE gel (Thermo Fisher Scientific) and proteins were transferred to a PVDF membrane and then immublotted with specific antibodies following blocking by 5% skim milk in 1 ×TBS (Tris buffered saline). IHC was performed by standard HRP-DAB staining protocol for paraffin-embedded tissues fixed in 10% neutral buffered formalin with a counter staining by haematoxylin and eosin. Following antibodies are used for protein detection in Western blot and immunohistochemistry: anti-estrogen receptor α (ERα) rabbit polyclonal antibody (MC-20) (Santa Cruz Biotechnology, # sc-542), rabbit polyclonal anti-SRSF3 antibody (Abcam, # ab125124), anti-SRSF1 mouse monoclonal antibody (clone 96) (Thermo Fisher scientific, # 32-4500), anti-HER2/ErbB2 rabbit monoclonal antibody (Cell Signaling Technology, # 2165) for mouse and rat c-neu/Erbb2, anti-Eif4a2 rabbit polyclonal antibody (Abcam, # ab31218), anti-β-tubulin antibody (Sigma, # T5201), anti-β-actin mouse monoclonal antibody (AC-15) (Santa Cruz Biotechnology, # sc-69879), and anti-GAPDH monoclonal antibody (Cell Signaling Technology, # 2118) for GAPDH.

### Serological analysis

Plasma samples from liver Srsf3-WT or Srsf3-KO mice of a year of age were subjected to serological assays to quantify plasma alanine transaminase (ALT), alkaline phosphatase, amylase activity, urea nitrogen and bilirubin by VetScan VS2 system (ABAXIS), or plasma estradiol concentration by ELISA, which was performed by Ani Lytics, Inc. (Gaithersburg, MD).

### Statistics

Log-rank test and Kaplan-Meyer plot were conducted using SigmaPlot (Systat Software Inc.). IHC images were analyzed by Aperio ImageScope (Leica Biosystems). The chi-squared test was used to compare the breast and liver tumor formation rates between Srsf3 WT and KO mice at 6 months. Gene expression levels across different groups, assessed through RT-qPCR, were evaluated using Student’s *t*-test. Biological differences with p < 0.05 were considered statistically significant.

## Results

### Srsf3 conditional KO in mouse mammary glands and liver tissues

Conditional KO of Srsf3 in mammary glands by MMTV-Cre and liver-specific KO of Srsf3 by Alb-Cre expression were based on Srsf3-floxed mice (19) (Figure S2A/B). We confirmed the specific homozygous KO of Srsf3 (Srsf3 KO) in mouse mammary gland tissues and liver tissues first by RT-PCR (Figure S2C-S2D) and subsequently by RNA-seq (Figure S2E-S2F), showing the KO of Srsf3 exon 2 and exon 3. Because the poison exon 4 is normally skipped in the WT mRNA (35,36), the detected SRSF3 mRNA having the Cre-mediated KO of exon 2 and exon 3 in breast and liver tissues appeared as the spliced product from the exon 1 to exon 5. The residual WT Srsf3 expression in the Srsf3 KO mammary gland tissues or liver tissues was detected and most likely originated from infiltrated leukocytes or other types of cells in the tissues lacking MMTV or Albumin promoter activities.

We noticed that mice with the conditional KO of Srsf3 in mammary glands and livers did not show embryonic lethality. However, the mice with liver conditional Srsf3 KO were born in a smaller size, less active and with less appetite. The most of them (∼90%) died by ∼14 days of birth (28) (Figure S3A). After chemical carcinogen diethylnitrosamine (DEN) injection of survived mice at day 15 of age, the death incidence of homozygous liver Srsf3 KO mice reduced to ∼24% in a month after birth (Figure S3B). However, the survived male and female mice from the death crisis recovered their body weight and size comparable to the Srsf3 WT mice by 2 months of age, consistent with previous study (30). Such growth failure or early death phenotype did not happen to the mice with mammary Srsf3 KO, nor to the mice bearing only heterozygous Srsf3 KO (hetKO) in the liver tissues (Figure S3B). Data suggest that conditional Srsf3 KO in liver had transient impacts on liver function responsible for the newborn growth, which subsequently could be overcome at the adulthood age. Serology liver function tests indicated that mice livers bearing Srsf3 KO displayed more damage to their liver function, with significant increase of ALT, ALP and Amylase activities, whereas urea nitrogen and total bilirubin were normal (Figure S4).

### Srsf3 KO prevents the Erbb2 breast cancer induction but enhances DEN-induced liver carcinogenesis

We then investigated the tissue-specific KO of Srsf3 on cancer induction in two well-established mouse cancer models: activated rat c-neu (V664E) replacement of mouse Erbb2-induced breast cancer (32) (Erbb2 breast cancer in this report), and DEN-induced liver cancer (33) (Figure 1A). In the DEN-induced mouse liver cancer model, we observed that ∼24% of the mice with liver-specific KO of Srsf3 died by one month-old (Figure S3B), whereas no early death was observed in the mice bearing WT Srsf3 or hetSrsf3 KO.

**Figure 1.**
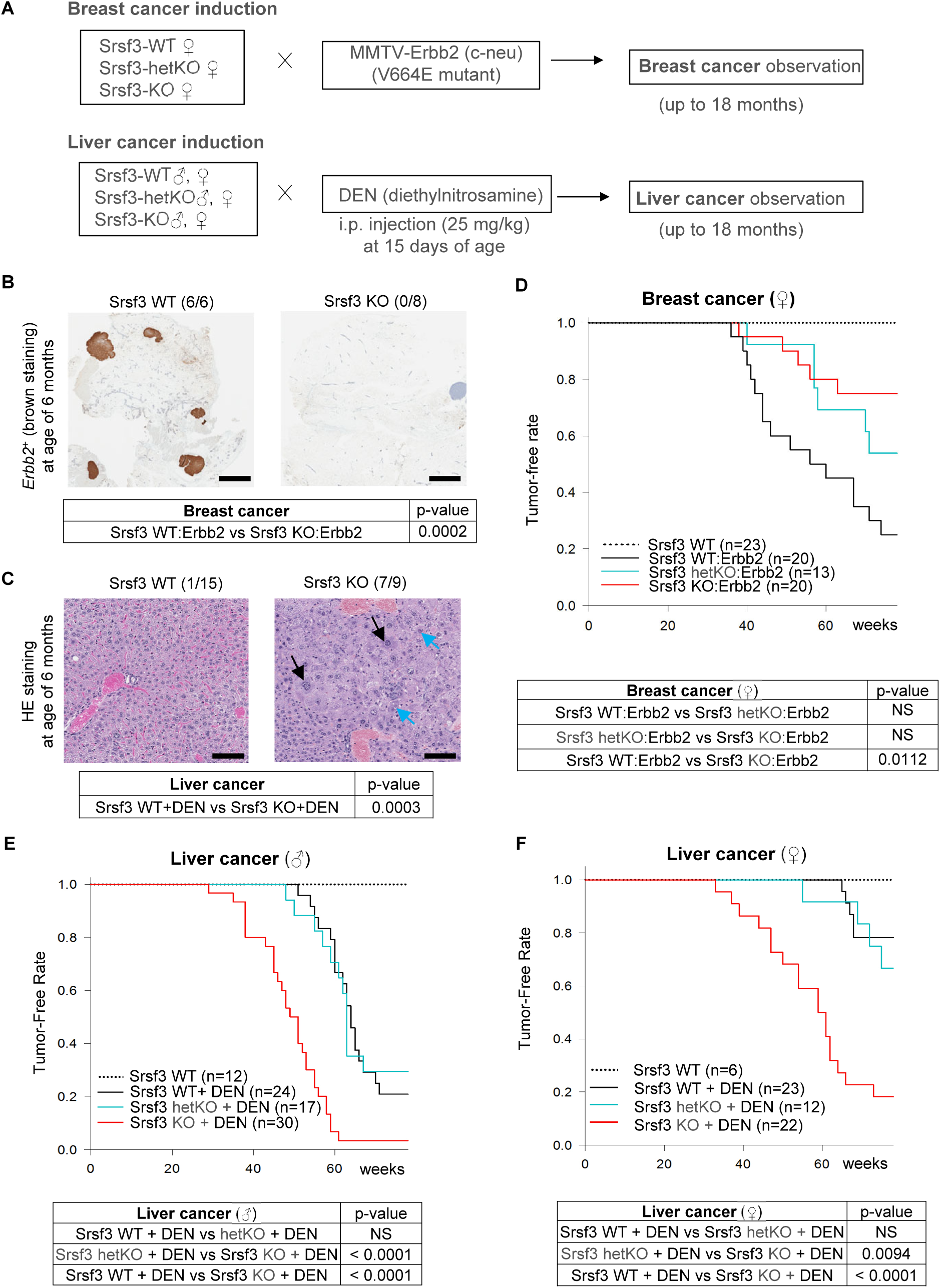
Srsf3-knockout (KO) delays development of Erbb2 breast cancer, but promotes liver cancer induction by DEN. (A) Flaw chart of mouse mammary-specific Srsf3 KO and Erbb2 breast cancer induction and liver-specific Srsf3 KO and liver cancer induction by DEN. Mammary cancer induction was assessed by crossing Srsf3-WT, hetKO (heterozygous knockout), KO female mice with male mice carrying a homozygous V664E mutant of rat c-neu/Erbb2 of which expression is under control by a MMTV promoter (MMTV). Liver cancer induction in mice with liver-specific Srsf3 knockout was performed by the intraperitoneal (i.p.) injection of diethylnitrosamine (DEN) at 25 mg/kg at 15 days of age. Tumor formation was surveyed by palpation weekly up to 18 months of age. (B) Erbb2 (c-neu) staining of Srsf3 WT and Srsf3 KO mammary tissues at six months of age. Six of six Srsf3 WT mice examined developed Erbb2-tumors, but none of all eight (0/8) Srsf3 KO mice had Erbb2 tumors. Scale bar, 2 mm. (C) H&E staining of Srsf3 WT and Srsf3 KO liver tissues at age of six months. Seven of nine (7/9) Srsf3 KO mice developed hepatocellular carcinoma (HCC), but only one of 15 Srsf3 WT mice developed HCC. Black arrow, karyomegaly. Blue arrow, intranuclear intracytoplasmic invagination. Scale bar, 100 µm. The p-values in (B) and (C) were calculated by Chi-squared test. (D, F) Kaplan-Meyer curves for the observed tumor formation rates were plotted till 18 months of age for Erbb2 breast cancer (D) and DEN-induced liver cancer (F) in female mice with or without mammary Srsf3 hetKO or homozygous Srsf3 KO. (E) DEN-induced liver tumor in male mice with or without liver heterozygous or homozygous Srsf3 KO. The p-values were determined by log-rank test between indicated groups.

The mice with conditional KO of Srsf3 in the mammary glands bearing the mutant rat c-neu and in the liver tissues with a peritoneal DEN injection (25 mg/kg at 15 days of age) were examined weekly by palpation for breast tumor development from a month of age and liver tumor development from 6 months of age, up to 18 months (a total of 78 weeks). Additionally, a subgroup of mice was sacrificed at the 6^th^ month for tissue pathological changes. The Kaplan-Meyer curves for animals with palpable tumors were plotted to 18 months. We discovered that Srsf3 plays opposite roles in the development of breast and liver cancers. In the Erbb2-induced breast tumor model, Srsf3 was oncogenic in promoting breast tumor formation because all mammary glands with WT Srsf3 expression in six Erbb2 mice exhibited c-neu (Erbb2)-positive tumor foci, but none of eight Erbb2 mice with the Srsf3 KO had any detectable tumor foci in the mammary glands at the 6^th^ month of animal age (Figure 1B). In contrast, we found seven out of nine mice with liver Srsf3 KO, but only one of 15 mice bearing WT Srsf3 with DEN injection, developed pathological hepatocellular carcinoma at the 6^th^ month of mice age (Figure 1C).

By the 18 months, Srsf3’s oncogenic role in the development of mouse breast tumor became more obvious. A significant delay (p = 0.0112) in tumor development in Erbb2 mice with Srsf3 KO was observed when compared to Erbb2 mice with WT Srsf3 (Figure 1D). The Erbb2 mice with hetSrsf3 KO showed intermediate outcome in overall tumor formation, and mice without Erbb2 failed in breast tumor induction during 18 months of observation period (Figure 1D). In contrast, Srsf3 was found to be tumor suppressive and played an important sex-disparity role in the development of liver tumor during 18 months of observation. We found that both male (Figure 1E) and female (Figure 1F) mice with Srsf3 KO show a much higher incidence in DEN-induced liver cancer formation (p < 0.0001) when compared to WT Srsf3 mice. The mice with hetSrsf3 KO showed no difference in liver cancer formation rate when compared to WT Srsf3 mice (Figure 1E and 1F), suggesting a single copy of Srsf3 allele expression (hetSrsf3 KO) is sufficient for delaying liver cancer formation. Because of female sex hormone, female mice (Figure 1F) in general are more resistant than male mice (Figure 1E) to liver cancer induction by DEN (p< 0.01)(33). Surprisingly, we found that Srsf3 KO diminished this sex disparity in the liver cancer development induced by DEN (compare Figure 1E with Figure 1F).

### Srsf3 regulates the expression of different sets of genes in association with Erbb2 breast cancer distinguishable from DEN-induced liver cancer

To understand how Srsf3 KO in mammary glands and liver tissues could have an opposite effect on carcinogenesis which might associated with altered gene expression and RNA splicing in two different tissues, we conducted RNA-Seq for the Erbb2-induced breast cancer tissues with Srsf3 KO and Srsf3 WT (Figure S5A-S5B, Table S1), and DEN-induced liver cancer tissues with Srsf3 KO and WT Srsf3 (Figure S5D-S5E, Table S2). We found that the gene expression profile from breast cancer affected by Srsf3 KO was very different from liver cancer affected by Srsf3 KO. This could be interpreted as different carcinogens were used, with Erbb2 as an oncogene in breast cancer induction and DEN as a chemical carcinogen in liver cancer induction, in addition to tissue-specific gene expression.

In the Erbb2 breast cancer, a total of 425 differentially expressed genes were identified, of which 159 genes were upregulated and 266 genes were downregulated in Srsf3 KO breast cancer when compared to WT Srsf3 breast cancer (Figure S5B, Table S1). The top enriched gene sets in upregulated pathways were involved in hypoxia, mitotic spindle, epithelial mesenchymal transition and p53 pathways, but the top enriched gene sets in the downregulated pathways were involved in interferon responses (Figure S5C). The heatmap showed the top 25 genes with abundantly upregulated or downregulated expression (at least one sample have RPKM≥2) (Figure S6A). By NanoString technology, we verified a differential expression of top 40 genes identified by RNA-seq (Figure S6B, Table S3). Among them, Pcbp4, Ahnak2 and Ctif are the genes involved in RNA processing and translation initiation, and Stat2, Il18bp and Casp7 are the genes involved in interferon response pathway. Interestingly, 20 out of the 40 NanoString-validated genes at the 6^th^ month of animal age (Figure S7A, Table S3) persistently exhibited the same upregulation or downregulation in Srsf3 KO breast tissues collected by 18 months when compared to WT Srsf3 breast tissues.

In DEN-induced liver cancer tissues, we found 1017 genes significantly upregulated and 927 genes significantly downregulated in Srsf3 KO versus WT Srsf3 liver cancers (Figure S5D/E, Table S2).The genes upregulated in Srsf3 KO liver tissues were involved in oxidative phosphorylation, epithelial mesenchymal transition, fatty acid metabolism, and E2F targets based on the GSEA hallmark gene-sets pathway analysis, whereas those downregulated genes in Srsf3 KO liver cancer were the genes responsible for interferon responses as observed in Srsf3 KO breast cancer, coagulation, and KRAS signaling (Figure S5F). Subsequently, we identified the top 25 each of upregulated and down-regulated genes in Srsf3 KO liver cancer tissues (Figure S6C) and verified the expression of top 15 upregulated and downregulated genes by NanoString analysis (Figure S6D, in red, Table S3). Besides these top regulated genes, we also validated the expression of genes with important functions (Figure S6D, in black). Nol3, Egfr and Ahnak2 are genes involving in RNA processing. Cxcl11, Zbp1 and Fgl2 are genes identified in the interferon pathway. Slc12a2 and Errfi1 are Srsf3-regulated genes identified in our previous publication (7). Interestingly at 6^th^ month of animal age, 11 out of the 43 validated genes exhibited the same up- or down-regulation in DEN-induced Srsf3 KO liver cancer when compared to DEN-induced Srsf3 WT liver cancer at the 18 months of observation (Figure S7B, Table S3).

We also analyzed the differential expression of Sox4, E2f1, Trpv4 and Trim6, which are the genes important for cancer stem cell behavior and function (37–42) (Figure 2). Although Srsf3 KO had no effect on the expression of Sox4, E2f1, Trpv4, and Trim6 in the Srsf3 KO Erbb2 breast cancer when compared to the Srsf3 WT Erbb2 breast cancer (Figure 2A-2D), we discovered that Srsf3 KO increased the expression of Sox4, E2f1, Trpv4 and Trim6 expression in DEN-induced liver cancer when compared to DEN-induced Srsf3 WT liver cancer (Figure 2E-2H). We further verified the increased expression of Sox4, E2f1, Trpv4, and Trim6 in DEN-induced, Srsf3 KO liver cancer tissues by qRT-PCR (Figure 2I-2L). Together, these data clearly indicate that Srsf3 functions differently in the tissue context by altering the expression of specific sets of genes which could partially explain the different roles Srsf3 might play in the Erbb2 breast cancer from DEN-induced liver cancer.

**Figure 2.**
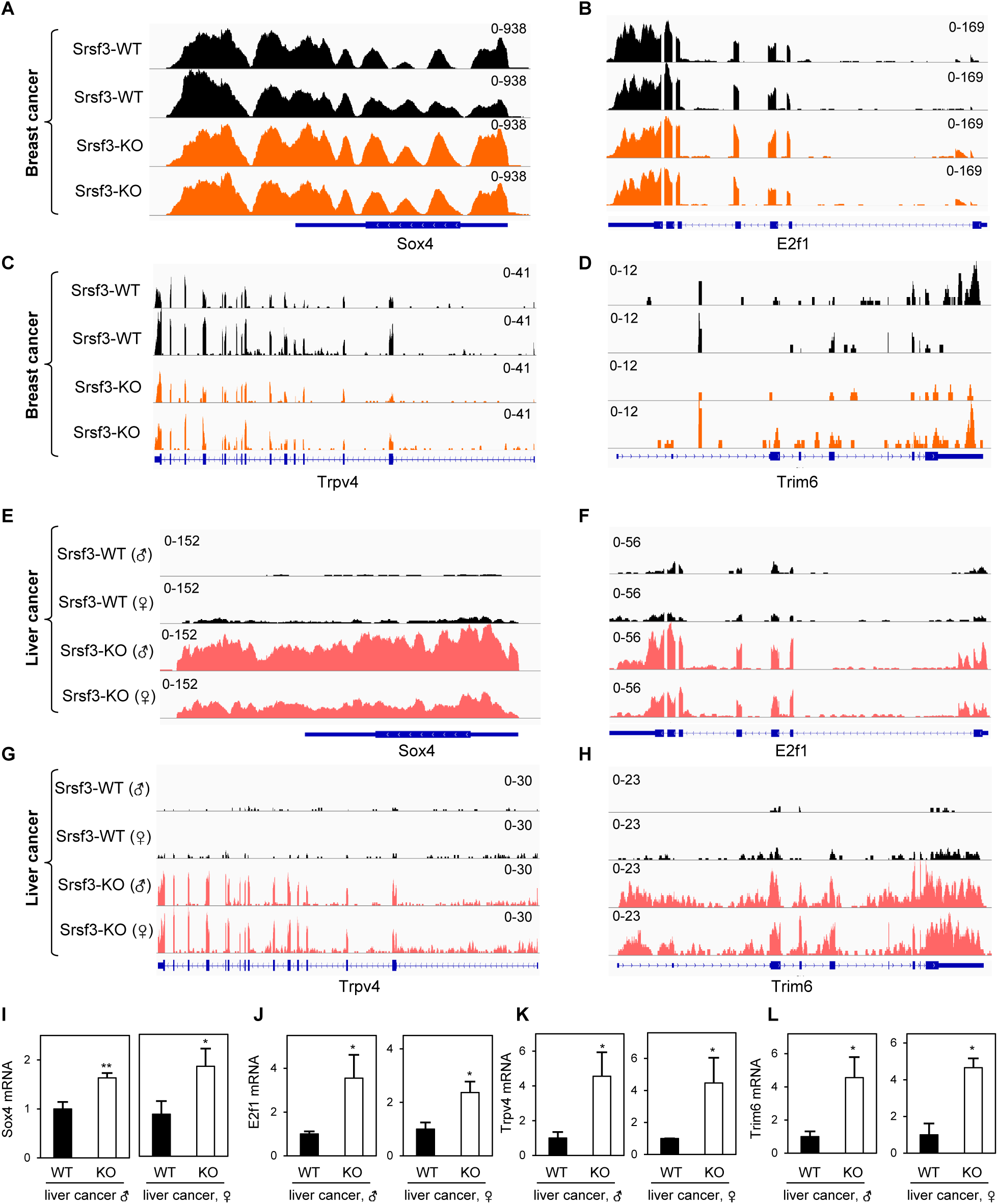
SRSF3 regulation of a subset of genes differentially from breast to liver cancer. The IGV illustrations show a distribution ofRNA-Seq reads mapped to Sox4, E2f1, Trpv4, and Trim6 genes in breast cancer samples (A-D) and liver cancer samples (E-H). The decreased expressions of Sox4, E2f1, Trpv4, and Trim6 genes in liver cancer tissues (n=3-5) were validated by TaqMan RT-qPCR (I-L). *, p < 0.05; **, p < 0.01 by Student’s *t*-test.

### Identification and validation of Srsf3-regulated splicing events in Erbb2 breast cancer and DEN-induced liver cancer

To understand Srsf3-induced changes in RNA splicing, we performed alternative splicing analysis using rMATS (34) and identified 1118 Srsf3-responsive splicing events in 899 genes in Erbb2 breast cancers with or without Srsf3 KO (Figure 3A, Table S4). We further selectively validated the Srsf3-regulated exon skipping of Slain2 exon 7/8, Fam221a exon 6, and Ralgapa1 exon 41, the selection of an alternative 5’ splice site in Mynn exon 2, Srsf5 exon 1, and Tmem161b exon 7, and the selection of an alternative 3’ splice site in Depdc5 exon 26 (Figure 3B-F, Figure S8A-B). Srsf3 KO in Erbb2 breast cancer was also found to promote Yip2 intron 9 retention and activate a cryptic intron within the exon 4 of Srsf1 (Figure S8C-D) to inhibit Srsf1 protein expression (Figure S9A)(7), consistent with the reported Srsf1 as protooncogene(43–46).

**Figure 3.**
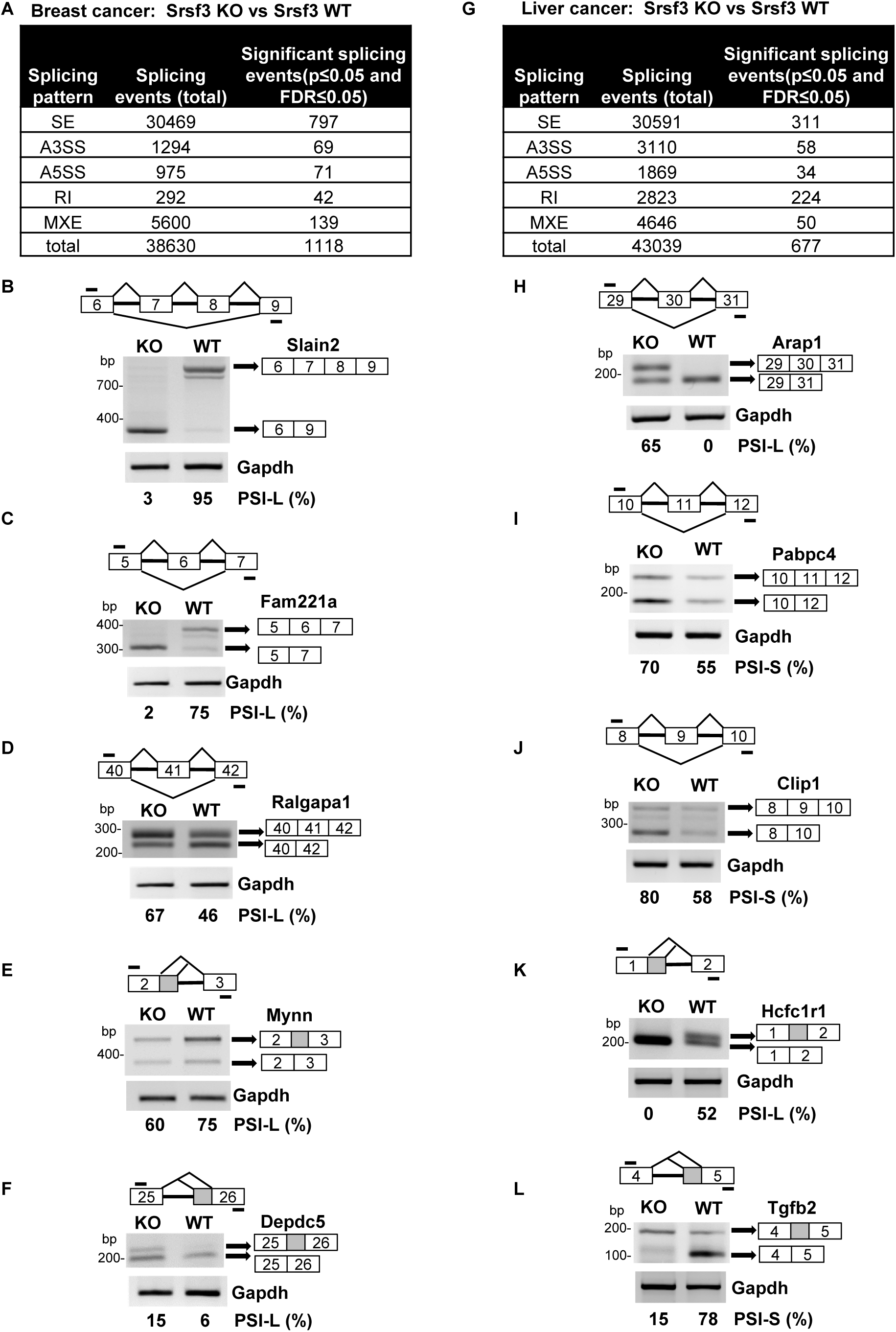
Identification and validation of differential RNA splicing events from Srsf3 KO and Srsf3 WT breast cancer and liver cancer.(A-F) Srsf3 KO regulates alternative RNA splicing events from breast Srsf3 WT cancer to Srsf3 KO cancer. (A) Genome-wide alternative RNA splicing events regulated by Srsf3 KO and identified by rMATS analysis (p≤0.05, FDR≤0.05) from breast Srsf3 WT to Srsf3 KO cancer are summarized. SE, skipped exon; A5SS, alternative 5ʹ splice site; A3SS, alternative 3ʹ splice site; RI, retained intron; MXE, mutually exclusive exons. (B-F) The genes highly susceptible to Srsf3 KO regulation in breast cancer tissues were selectively verified by RT-PCR on exon skipping of the Slain2 exons 7-8 (B), Fam221a exon 6 (C), and Ralgapa1 exon 41 (C) and alternative 5ʹss splicing of Mynn exon 2 (E), and alternative 3ʹss splicing of Depdc5 exon 25. The primers used in RT-PCR are shown as bars above (forward primers) and below (reverse primers) each RNA diagram. Gapdh served as a loading control. PSI, percent spliced-in inclusion (L=long RNA isoform) or exclusion (S=short RNA isoform) of an alternative exon, intron, or splice site (% spliced-in = inclusion / sum of inclusion + exclusion for long RNA isoform or exclusion / sum of exclusion + inclusion for short RNA isoform). (G-L) Srsf3 KO regulates alternative RNA splicing events from liver Srsf3 WT cancer to Srsf3 KO cancer. (G) Genome-wide alternative splicing events regulated by Srsf3 KO and identified by rMATS analysis (p≤0.05, FDR≤0.05) from liver Srsf3 WT to Srsf3 KO cancer are summarized. (H-L) The genes highly susceptible to Srsf3 KO regulation in liver cancer tissues were selectively verified by RT-PCR on exon skipping of the Arap1 exon 30 (H), Pabpc4 exon 11 (I), and Clip1 exon 9 (J) and alternative 5ʹss splicing of Hcfc1r1 exon 1 (K) and alternative 3ʹss splicing of Tgfb2 exon 5 (L). See other details in (A-F).

In DEN-induced liver cancer, a total of 677 Srsf3-regulated splicing events of 575 genes were detected by rMATS (Figure 3G, Table S5). We selectively validated the exon skipping of Arap1 exon 30, Pabpc4 exon 11, Clip 1 exon 9, and Nprl3 exon 3, the selection of an alternative 5’ splice site of Hcfc1r1 exon 1 and Zfp512b exon 8, the selection of an alternative 3’ splice site of Tgfb2 exon 5 and Arv1 exon 3, and Kin intron 2 retention (Figure 3 H-L, Figure S8E-H). The exon skipping of Pabpc4 targeted by Srsf3 was reported in our previous study (5).

### Common Srsf3-responsive genes identified from breast cancer and liver cancer

Although the difference in Srsf3 KO-induced gene expression profiles identified above were notable and most likely resulted from the different organs (breast vs liver) and different carcinogens used in the tumor induction, we wanted to investigate the common Srsf3-responsive genes and to explore how Srsf3 plays its opposite roles in the development of Erbb2 breast cancer and DEN-induced liver cancer. By Venn diagram analysis (Figure 4A), we identified 41 genes commonly responsive to Srsf3 KO, with 15 genes significantly downregulated in breast cancer but upregulated in liver cancer, and 9 genes significantly upregulated in breast cancer but downregulated in liver cancer upon Srsf3 KO (Figure 4B)., The remaining 17 genes were common Srsf3 KO responders, of which 4 genes were upregulated and 13 genes were downregulated, including Srsf1, by Srsf3 KO in both Erbb2 breast cancer and DEN-induced liver cancer (Figure 4A). Srsf3 KO was found to reduce Srsf1 protein expression both from the Erbb2 breast cancer and DEN-induced liver cancer (Figure S9A-9D). Major Facilitator Superfamily Domain Containing 4A protein (Mfsd4a) involved in glucose transmembrane transport is on the top list of dysregulated genes, and is down-regulated 16-fold in Srsf3 KO Erbb2 breast cancer, but up-regulated 293-fold in DEN-induced Srsf3 KO liver cancer (Figure 4B). This opposite response of Mfsd4a to Srsf3 KO from Erbb2 breast cancer to DEN-induced liver cancer was remarkable by IGV visualization of RNA-seq reads-coverage (Figure 4C) and easily validated by NanoString analysis (Figure S6B and S6D). Similarly, Htatip2 (Tip30), a tumor suppressor in various cancers (47–49), was downregulated in Erbb2 breast cancer, but upregulated in DEN-induced liver cancer following Srsf3 knockout (Figure S10A). Srsf3 KD in Hepa 1-6 cells also promoted the expression of Htatip2 (Figure S10B-C).

**Figure 4.**
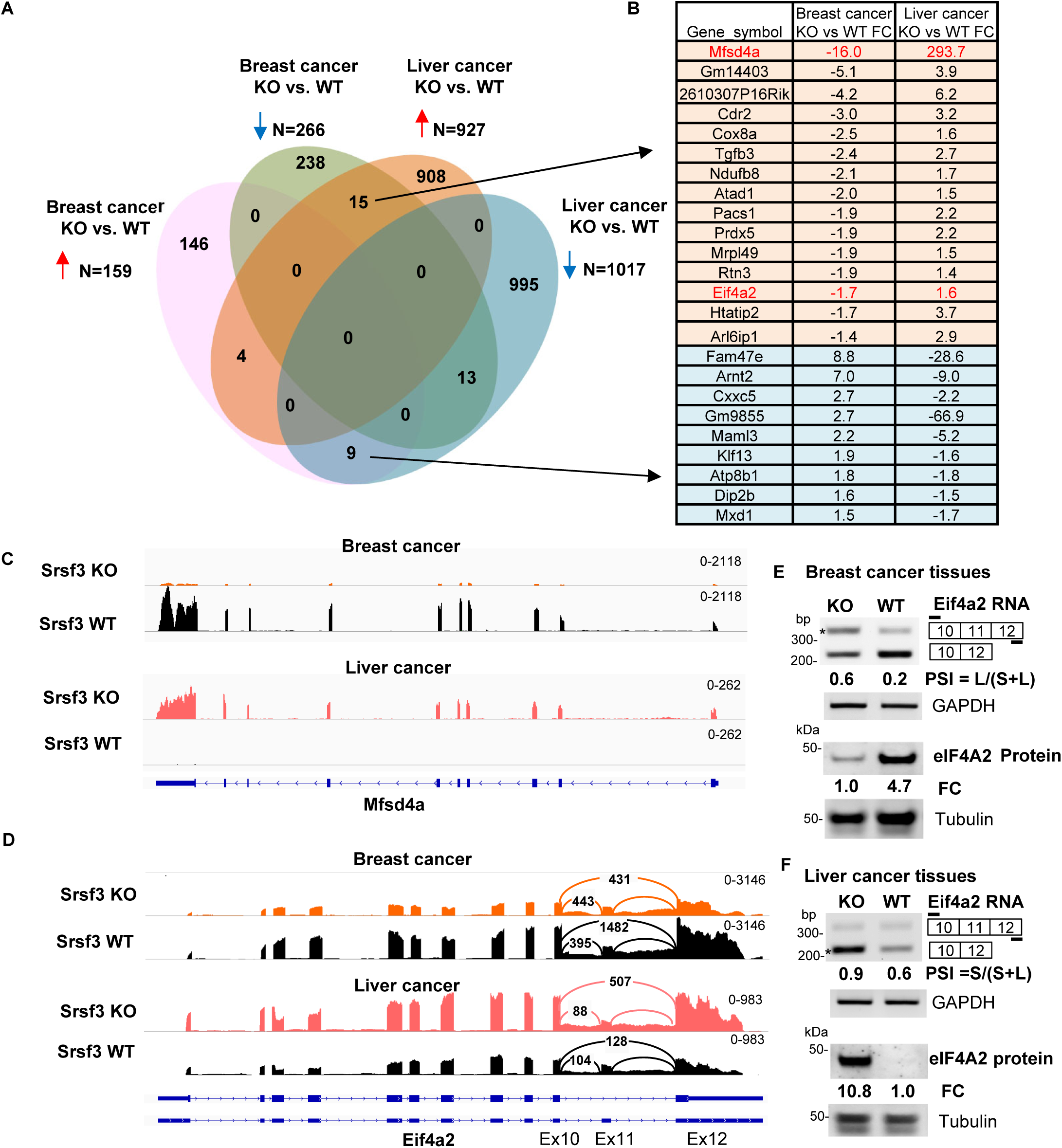
Differential expression and RNA splicing of the identified Srsf3 targets in response to Srsf3 KO from breast cancer to liver cancer. (A) The Venn diagram shows the overlapped genes with differential expression changes (adjusted p ≤ 0.05) from breast cancer to liver cancer by RNA-seq analysis. (B) RNA-seq analyses identified 15 overlapped genes susceptible to Srsf3 KO significantly down-regulated in breast cancer but upregulated in liver cancer, and 9 overlapped genes susceptible to Srsf3 KO significantly upregulated in breast cancer but down-regulated in liver cancer. Eif4a2 in red is the only gene undergoing alternative RNA splicing by Srsf3 KO in both breast and liver cancer tissues detected by rMATs. (C) Srsf3 KO-induced differential expression of Mfsd4a from breast cancer tissues to liver cancer tissues identified by RNA-seq reads-coverage. Srsf3 KO blocks Mfsd4a expression in breast cancer but induces Mfsd4a expression in liver cancer when compared to its Srsf3 WT cancer tissues. (D) Srsf3 KO-induced differential alternative RNA splicing of Eif4a2 from breast to liver cancer tissues identified by RNA-seq reads-coverage. The numbered arches represent splice junction reads detected in Sashimi plots. (E and F) Alternative splicing validation by RT-PCR with an indicated primer pair (short lines) shown above the diagram of alternatively spliced exons (numbered boxes). Below is the Western blot analysis of Eif4A2 protein expression. Srsf3 KO promoted the exon 11 inclusion of Eif4a2 splicing and thereby inhibited Eif4a2 protein expression in breast cancer (E) but reduced the exon 11 inclusion and increases Eif4a2 protein expression in liver cancer when compared to its Srsf3 WT cancer tissues (F). Eif4a2 exon 11 inclusion and exclusion in Srsf3 KO and WT breast and liver cancer tissues were verified by RT-PCR with GAPDH RNA served as a loading control (E and F). PSI, percent spliced-in of the alternative exon(s); large size band (E)(% inclusion = inclusion/sum of inclusion + exclusion); small size band (F) (% exclusion = exclusion/sum of inclusion + exclusion); FC (fold change); comparison between the sum of both inclusion and exclusion isoform in WT and KO tissue, normalized to Gapdh in RT-PCR analysis. In Western blot analysis, the FC of Eif4a2 between KO and WT cancer tissues was calculated after normalizing to β-tubulin.

Eukaryotic translation initiation factor 4A2 (Eif4a2 or DDX2B) was identified as another Srsf3 KO responder. Eif4a2 gene has 12 exons and its exon 11 is a “poison exon”. Inclusion of the exon 11 in Eif4a2 RNA leads to Eif4a2 RNA degradation by non-sense mediated decay. Only the Eif4a2 mRNAs with the exon 11 exclusion can encode a full-length Eif4a2 protein. Therefore, inclusion of the exon 11 in Eif4a2 RNA splicing will reduce the expression of full-length Eif4a2 protein. We found by RNA-seq that Srsf3 KO promoted the exon 11 inclusion of Eif4a2 splicing and reduced Eif4a2 RNA level due to induction of Eif4a2 RNA degradation in breast cancer (Figure 4D). In contrast, Srsf3 KO in liver tumor cancer promoted the exon 11 exclusion and enhanced the expression of Eif4a2 RNA (Figure 4D). RT-PCR analyses of the total RNA from Erbb2 breast cancer and DEN-induced liver cancer tissues and Western blot analyses of the tissue lysates from the Erbb2 breast cancer and DEN-induced liver cancer tissues, respectively, confirmed Srsf3 KO increasing the exon 11 inclusion of Eif4a2 mRNA and decreasing the eIF4A2 protein production in the Erbb2 breast cancer (Figure 4E), but promoting the exon 11 exclusion of Eif4a2 and increase the production of eIF4A2 protein in the DEN-induced liver cancer (Figure 4F).

Consistently with Erbb2 cancer tissues, knockdown of Srsf3 expression in mouse Erbb2^+^ NF639 cells and two human breast cancer cell lines ER^+^ MCF7 and HER2^+^ SKBR3 cells was also found to promote exon 11 inclusion of Eif4a2 (Figure S10D) and subsequently to inhibit the production of eIF4A2 protein (Figure S10E).

### Srsf3 KO-mediated reduction of ERα and Foxa might lead to loss of sex disparity in the development of DEN-induced Srsf3 KO liver cancer

Liver cancer is one of six most frequently diagnosed cancer worldwide, and men show significant higher liver cancer incidence and death over the women (25). Male rodents are also prone to liver cancer when compared to females by chemically induced carcinogenesis (33,50). Estrogen and estrogen receptor ERα are protective for female from liver cancer (33,51,52). Foxa gene family members are contributors to sex disparity in liver cancer and female Foxa1/2-deficient livers are prone to HCC as male livers. ERα and Foxa gene family coregulate the certain genes responsible for female mouse resistance to liver cancer induction by DEN (53). Consistent with the previous report, our result also showed higher DEN-induced liver cancer incidence in Srsf3 WT male mice (Figure1E) than in Srsf3 WT female mice (Figure 1F). Surprisingly, we found that Srsf3 knockout reduced the gender disparity of DEN-induced liver cancer incidence between male and female mice (compare Figure 1E to Figure 1F, Fig S11). Given the above observations and to seek for how Srsf3 KO led to a reduction of gender disparity in DEN-induced liver cancer, we examined potential effect of Srsf3 KO on estrogen signaling (33,51,52) and Foxa gene family (53). RNA-seq analyses showed that DEN-induced Srsf3 KO liver cancer in both female and male mice exhibited a significantly reduced expression of Foxa1/2/3 and ERα than DEN-induced Srsf3 WT liver cancer (Figure 5A), which could be verified by quantitative RT-PCR (Figure 5B, Figure S12A). Western blot analysis and immunostaining of DEN-induced liver cancer tissues showed the decreased level of ERα protein in the Srsf3 KO female and male livers (Figure 5C, Figure S9C). Interestingly, the Srsf3 KO-mediated reduction of ERα protein in liver tissues led to an increased level of plasma estradiol E2 in Srsf3 KO female mice than the corresponding Srsf3 WT female mice (Figure 5D), suggesting impaired uptake of estradiol E2 by hepatocytes in the Srsf3 KO female mice.

**Figure 5.**
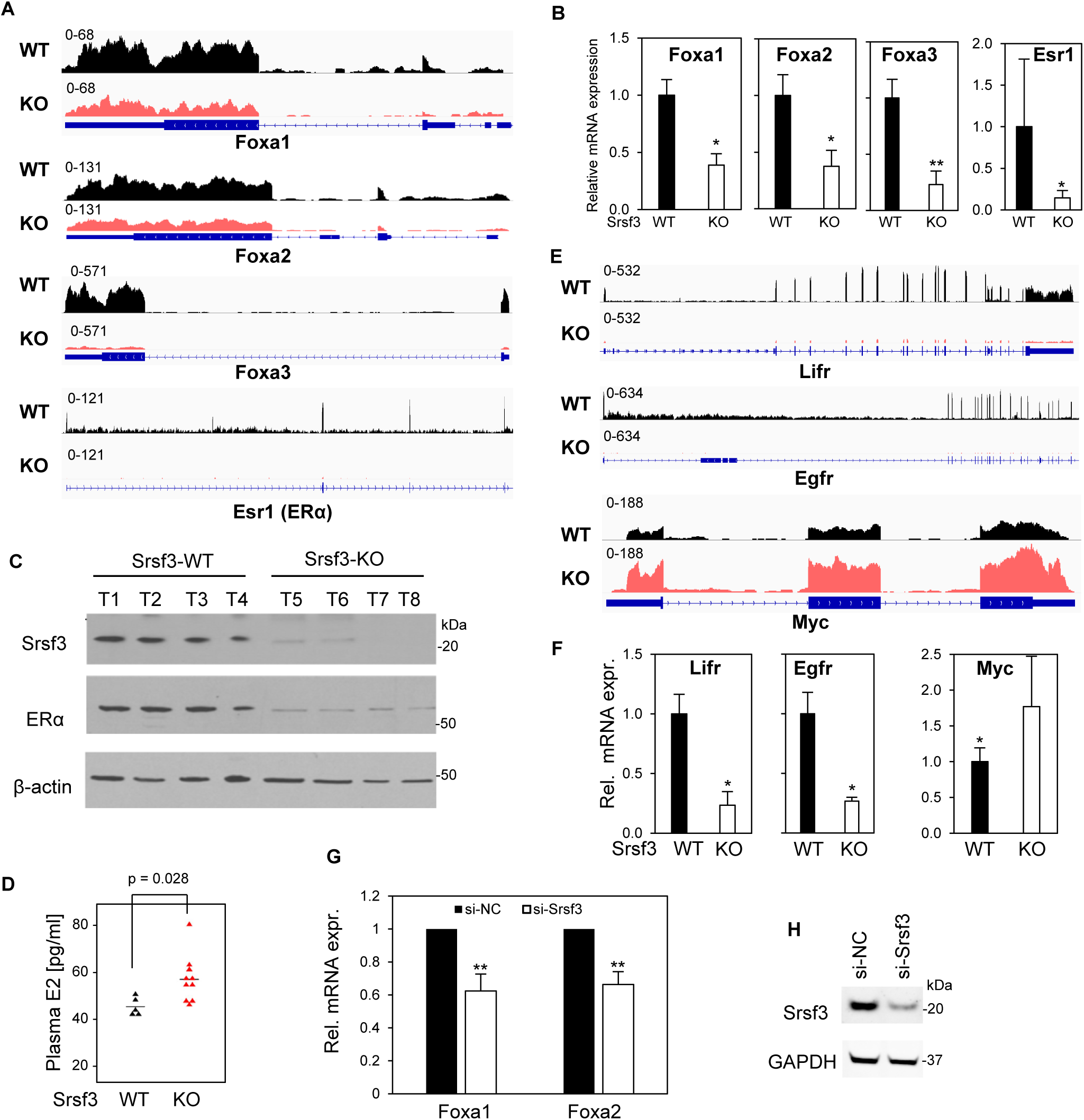
Srsf3 KO in female livers reduces the sex disparity in development of liver cancer by suppressing the expression of estrogen receptor ERα, Foxa gene family, Lifr, and Egfr, but increasing Myc expression. (A and E) Srsf3 KO led to decreased expression of Foxa1, Foxa2, Foxa3, Esr1(ERα), Lifr, and Egfr but increased expression of Myc. RNA-seq reads-coverage of indicated genes in female Srsf3-WT (black) and Srsf3-KO liver tumor (red) were visualized by IGV. (B and F) Validation of RNA expression for Foxa1,2,3 and Erα (B) and Lifr, Egfr, and Myc (F) in female Srsf3-WT and Srsf3-KO liver cancer tissues by RT-qPCR. Five samples in each group were examined. (C) Western blot examination of SRSF3 and ERα protein expression in female liver tumors. The β-actin served as loading control. (D) ELISA assay for plasma estradiol (E2) levels in Srsf3-WT and Srsf3-KO mice. (G and H) Knockdown of Srsf3 expression in a mouse liver cancer cell line Hepa1-6 reduced the expression of Foxa1 and Foxa2. Hepa1-6 cells were transfected with 40 nM of siRNA and total RNA and proteins were harvested at 48h after transfection. Foxa1 and Foxa2 expression levels were measured by RT-qPCR (G). The data were averaged from four independent experiments. Srsf3 knockdown efficiency was examined by Western blot with Gapdh serving as a sample loading control (H). *, p < 0.05; **, p< 0.01 by Student’s *t*-test (B, D, F, and G).

Lifr, Egfr and Myc are three downstream genes coregulated by Erα and Foxa (53). Our RNA-seq and RT-qPCR analyses of DEN-induced Srsf3 WT and Srsf3 KO liver cancer show significant reduction of Lifr and Egfr expression, but increased expression of oncogenic Myc (54) in DEN-induced Srsf3 KO liver cancer in both male and female mice (Figure 5E and 5F, Figure S12B). We also confirmed the reduction of Foxa1/2 expression after Srsf3 was knocked down in mouse liver cancer cell line Hepa1-6 cells (Figure 5G/H).

## Discussion

As an important splicing factor, SRSF3 has profound functions in health and its aberrant expression has been associated with progression of many diseases and cancer (4). In this report, we demonstrated that SRSF3 at its physiological level functions as a tumor-promotor in development of Erbb2 breast cancer, but a tumor-suppressor in DEN-induced liver cancer, through regulation of RNA transcription or alternative splicing of a different set of genes. This observation adds SRSF3 as another gene that functions in a tissue- or cell type-dependent manner for its opposite oncogenic potentials, as reported for TGF beta (55,56), NF-kB (57), Pten (58,59), PML (60,61), BRD4 (62), etc.

SRSF3 was initially discovered as a proto-oncogene responsible for cell proliferation overexpressed in various types of cancers for cancer cell proliferation and tumor maintenance. The increased expression of SRSF3 in breast tumors tissues was found in correlation with breast cancer progression and 5-year overall survival (20). SRSF3, as a splicing factor, is responsible for 23% of alternative RNA splicing events identified in 682 invasive breast ductal carcinoma patients and higher expression of SRSF3 was linked with shorter survival time and poorer prognosis (63). Consistently, we found conditional Srsf3 KO in mouse mammary glands prevent c-neu (Erbb2) expression and thus, Erbb2 breast cancer induction, providing for the first time the direct *in vivo* evidence of Srsf3’s oncogenic feature. RNA-seq profiling showed that Srsf3 regulated the expression of 425 genes including Mfsd4a, Pcbp4, Ahnak2 and Ctif, and 1118 RNA splicing events in 899 genes, including exon exclusion of Slain2, Fam221a, and Ralgapa1, intron splicing of Yip2, and alternative 5’ or 3’ usage of Mynn, Depdc5, and Tmem161b. Altogether, the altered gene expression and alternative RNA splicing by Srsf3 KO in mouse mammary glands may contribute to Srsf3 KO prevention of Erbb2 breast cancer.

However, Srsf3 was found to be tumor suppressive in DEN-induced liver cancer. We found that Srsf3 KO in liver could promote the development of DEN-induced liver cancer. This observation is consistent with the report showing Srsf3 KO enhancement of liver cancer development with mouse aging (30). Other reports also suggest that Srsf3 at a physiological level is critical for hepatocyte maturation and metabolic function (28) and prevention of DNA damage (64). Interestingly, despite its tumor suppressive function, SRSF3 mRNA is highly expressed in HCC patients (65). Using HCC tissue array assays we also found an increased SRSF3 protein expression in 110 samples of HCC (grade 2 and 3) compared to 10 samples of normal liver tissues (Figure S1), as seen in many other cancer tissues (4,7,20). The increased expression of SRSF3 in cancer cells is essential for cancer cell proliferation (20) and loss of SRSF3 expression leads to cell apoptosis (20) and senescence (21), and genome instability (64). The increased expression of tumor-suppressors (66) is common in many cancer types, including cyclin dependent kinase inhibitor p16Ink4a in glioma and HPV-induced cancers (67–69), CDK4/6 inhibitor p18Ink4c in cervical cancer and glioblastoma (70,71), and CDK2 and PCNA inhibitor p21 in many types of cancer (72).

We found that in the liver, Srsf3 regulated the expression of 1017 genes and 677 splicing events in 575 genes, including the expression of Hnf1α, Hmgcr, Srebp1, Srebp2, and Scap, and alternative RNA splicing of Fn1, Myo1b, Insr, which were previously reported (28–30), Srsf3 KO in mouse hepatocytes, but not in mouse mammary glands, promoted the expression of Sox4, E2f1, Trpv4, and Trim6, and alternative RNA splicing of Arap1, Pabpc4, Clip1, Hcfc1r1, Tgfb2, and Kin. Moreover, we identified that 24 out of 41 genes (Figure 4B) responsive to Srsf3 KO exhibited opposite expression profiles between breast cancer and liver cancer. In particular, Srsf3 KO in mouse hepatocytes increased the expression of Mfsd4a (73) and Eif4a2 (74), but Srsf3 KO in mouse mammary glands decreased the expression of these genes. Others, including Cox8a, Ndufb8, and Atad1, which are the genes involved in oxidative phosphorylation pathway (29), also exhibited an opposite regulation by Srsf3 KO in our Erbb2 breast cancer and DEN-induced liver cancer. These gene expression patterns exemplify how Srsf3 could act as a proto-oncogene in Erbb2 breast cancer, but a tumor suppressor in DEN-induced liver cancer. However, the molecular mechanism of how Srsf3 plays an opposite oncogenic function in breast and liver remains to be actively investigated.

As one of the six most frequently diagnosed cancers worldwide, liver cancer has an incidence and mortality rate that is twice as high in men compared to women (25). Male mice are prone to liver cancer compared to females when subjected to DEN-induced carcinogenesis (33) (Figure S11A). Surprisingly, Srsf3 knockout decreases the gender disparity in the development of liver cancer (Figure S11B). The gender disparity in the DEN-induced liver cancer has been associated with high expression of DEN-induced serum IL-6 in males (33). Estrogen inhibits IL-6 secretion from liver Kupffer cells and reduces liver cancer risk in females, whose ERα loss also leads to liver cancer (33). Loss of Foxa1 and Foxa2 expression in female liver affects ERα binding to its targeted genes and thus increases DEN-induced liver cancer in female mice (53). Surprisingly, we found Srsf3 also contributes the gender disparity in liver cancer. Srsf3 KO in our study decreased the expression of ERα and Foxa1-3 and its downstream genes Lifr and Egfr but increased Myc expression both in female and male livers, thereby promoting liver cancer formation by disrupting sex disparity. Moving forward in high reverence is to investigate how Srsf3 regulates the expression of ERα as a target for understanding liver carcinogenesis and for development of potential new liver cancer therapies.

## Data availability

Raw data and analyzed RNA-seq data supporting the findings in this study have been deposited in the NCBI GEO database under accession numbers GSE276011.

## Acknowledgements

This work was supported by the Intramural Research Program of the National Institutes of Health, the National Cancer Institute, and the Center for Cancer Research (ZIASC010357 to Z.M.Z.). We thank Hassan Jumaa of Albert-Ludwigs Universitat, Freiburg, Germany, for providing us the Srsf3^flox/flox^ mice. Rui-Hong Wang of NIDDK for helping us to establish albumin promoter-mediated KO of Srsf3 in hepatocytes. Vincent Homman and Haibin Liu for mouse genotyping, Elijah Edmondson for tissue pathology.

## Author contribution

L.Y, M.A., and Z.M.Z. designed the study, performed all experiments and data analyses. L.Y. and M.A. performed all the animal experiments. A.L. and M.C. performed the bioinformatic analysis. V.M. performed RNA-seq analysis and interpretations. D.G. carried out mouse genotyping and interpretations. B.K. performed histological diagnosis. C.X.D. provided transgenic mice. D.R.L. designed early stage of animal study. N.J.G.W. provided Srsf3 transgenic mice. L.Y., M.A. and Z.M.Z. drafted the manuscript and finalized with input from all co-authors.

## Declaration of interests

The authors declare no competing interests.

## Figure legend

**Fig. S1.**
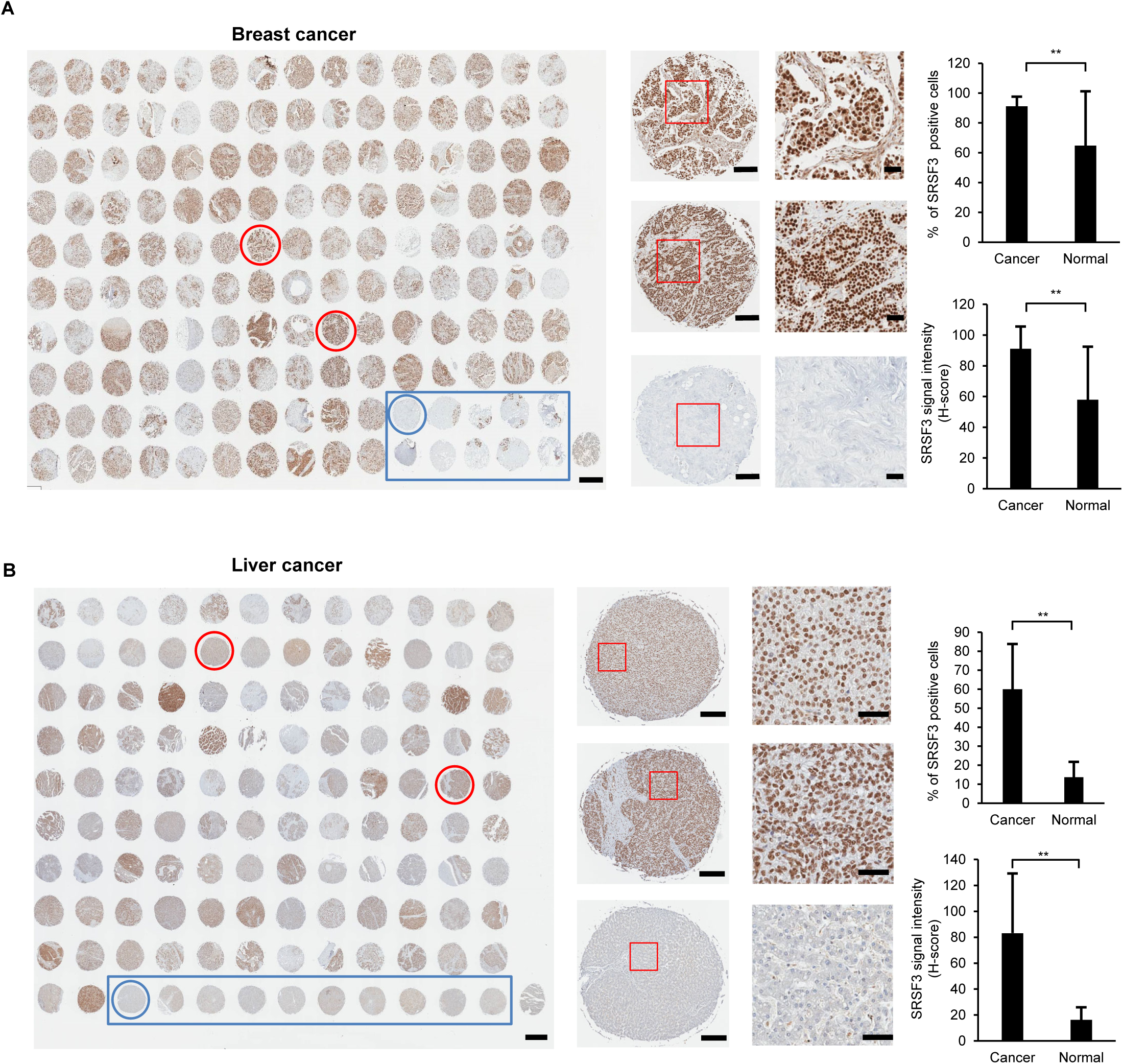
SRSF3 overexpressed in both human breast cancer and liver cancer tissues. (A) SRSF3 immunohistology staining of 70 paired human breast cancer tissues and 5 paired normal tissues (tissue array information could be found in https://www.tissuearray.com/BR1504b). (B) SRSF3 immunohistology staining of 110 unpaired, single human liver cancer tissues and 10 normal tissues (tissue array information could be found in https://www.tissuearray.com/tissue-arrays/Liver/BC03119b). For both panel A and panel B, Scale bars from left to right, 1mm, 200μm, 50μm. Normal tissues were highlighted in blue rectangle. SRSF3 signal was quantified by HALO software (https://indicalab.com/halo/). Statistical comparisons were computed by unpaired, two-tailed Student’s t test. ** p<0.01.

**Fig. S2.**
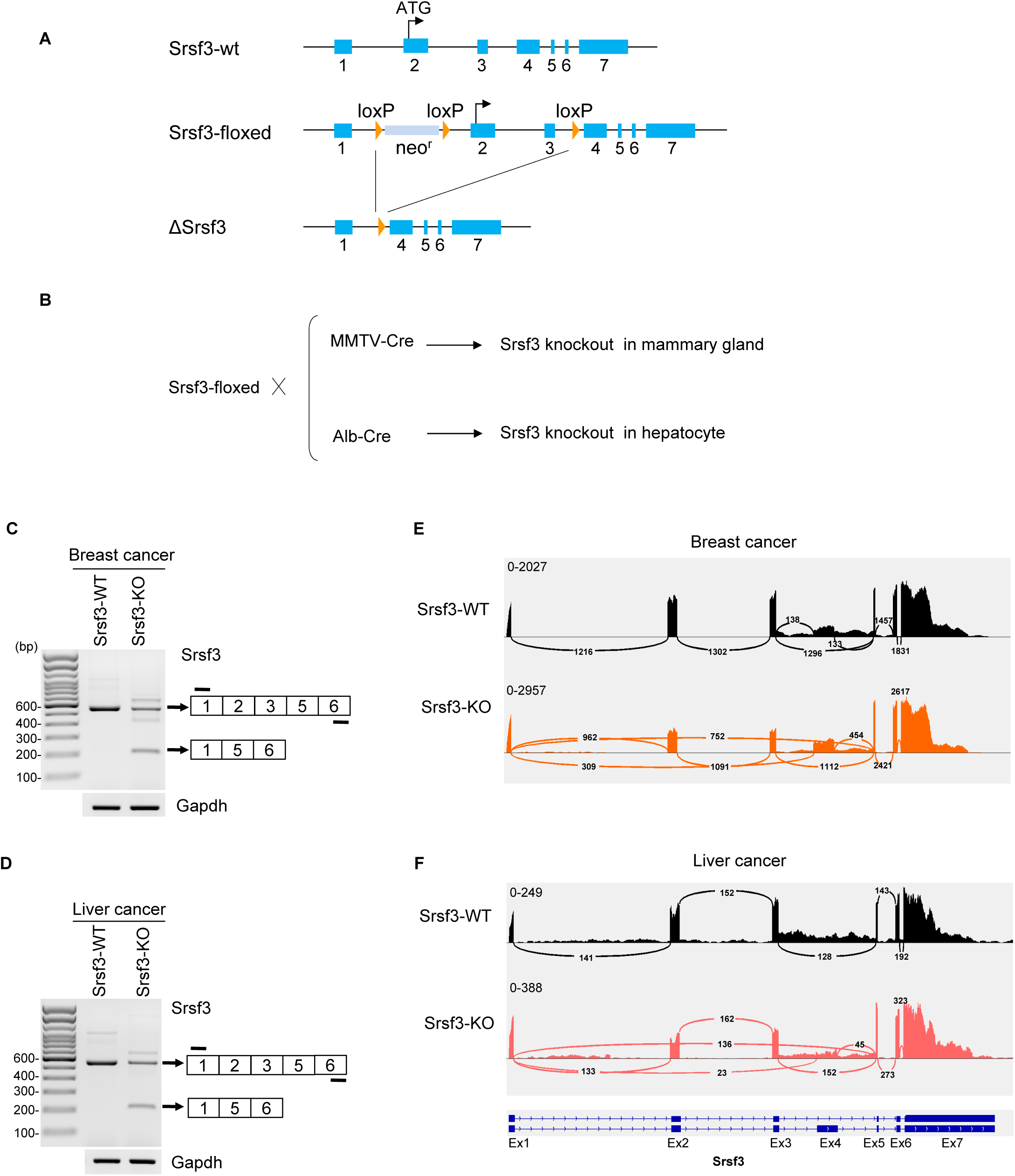
Strategies implemented for tissue-specific conditional knock-out (KO) of mouse Srsf3 in breast and liver tissues. (A) Diagrams of Srsf3 gene structure and knockout strategy by Cre-loxP technology. Srsf3 gene consists of seven exons with the start codon AUG in the second exon. In the Srsf3-floxed allele (Srsf3-floxed) as described (Jumaa H., et al. Current Biol. 9: 899-902, 1999), loxP sequences are inserted in the first and third introns, leading to excise the Srsf3 exon 2 and 3 from genomic DNA when Cre recombinase is expressed in cells to induce Srsf3 knockout due to loss of start codon (ΔSrsf3). Floxed NeoR, neomycin-resistant gene in the SRSF3 intron 1 under control by LoxP for cell selection. (B) Tissue-specific Srsf3 knockout is achieved by crossing Srsf3-floxed mice with Cre recombinase-expressing mice under a murine mammary tumor virus promoter (MMTV-Cre) or an albumin promoter (Alb-Cre), selectively active in mammary gland tissues or hepatocytes, respectively.(C and D) Sashimi plots showing the alternative splicing of Srsf3 gene in representative Srsf3 WT and KO breast cancer (C) and liver cancer tissues (D). The numbered arches represent splice junction reads detected from IGV. (E and F) RT-PCR validation for Srsf3 KO in breast cancer (E) and liver cancer tissues (F). Cre-LoxP recombination or KO of Srsf3 exons 2 + 3 mediated by tissue-specific Cre was confirmed by indicated primers from exon 1 and exon 6 (dash lines). GAPDH RNA served as a loading control.

**Fig. S3.**
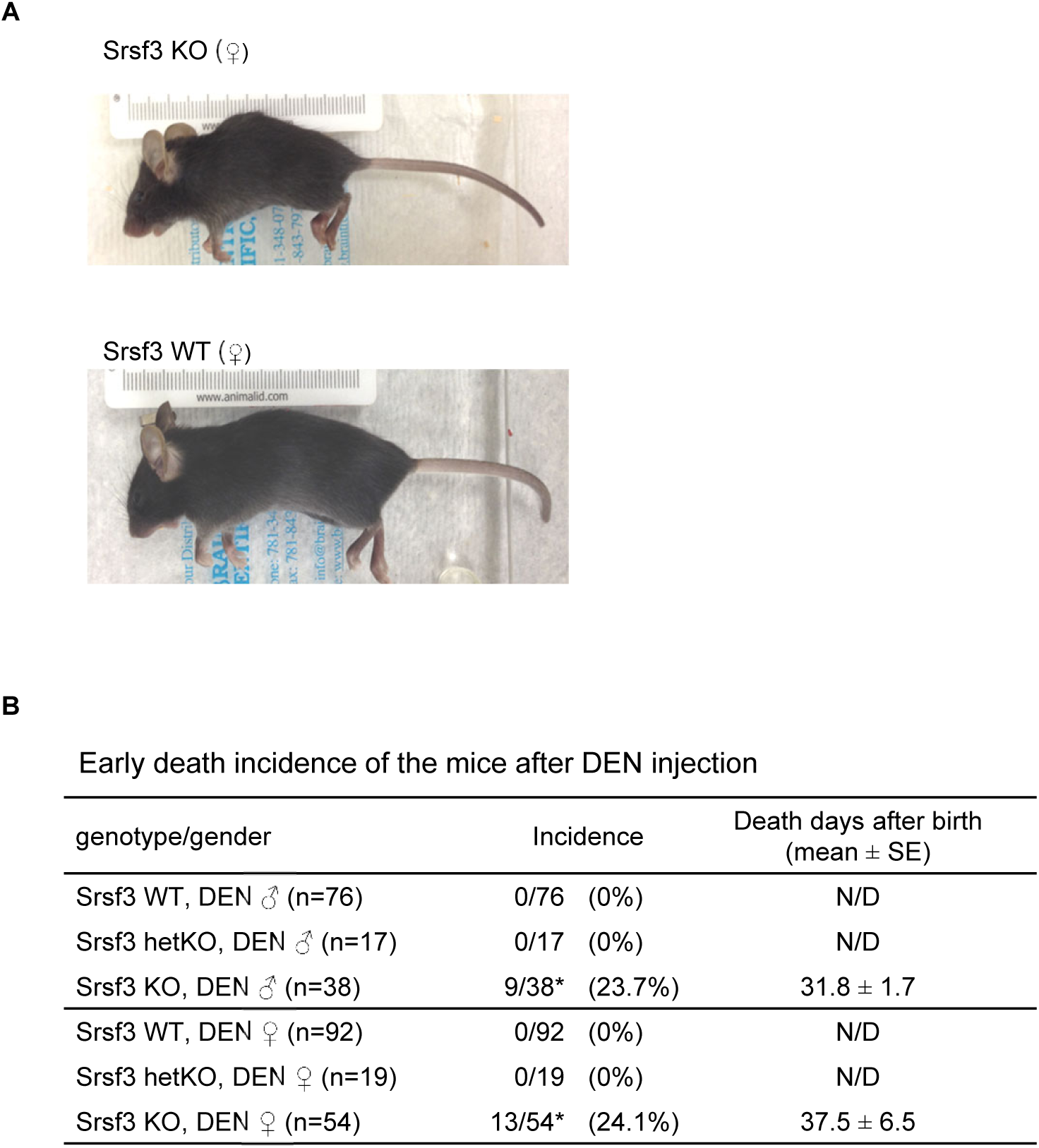
Summary of early phenotypes of liver Srsf3-KO mic. (A) Pictures represent typical liver Srsf3-KO and WT female mice at 2 weeks after birth. (B) Early death incidence of Srsf3-WT, -hetKO, and -KO mice by days after one dose DEN injection on day 15 of age. *Some of the survived mice with DEN injection were sacrificed for serology and pathology examination in the middle term of the study.

**Fig. S4.**
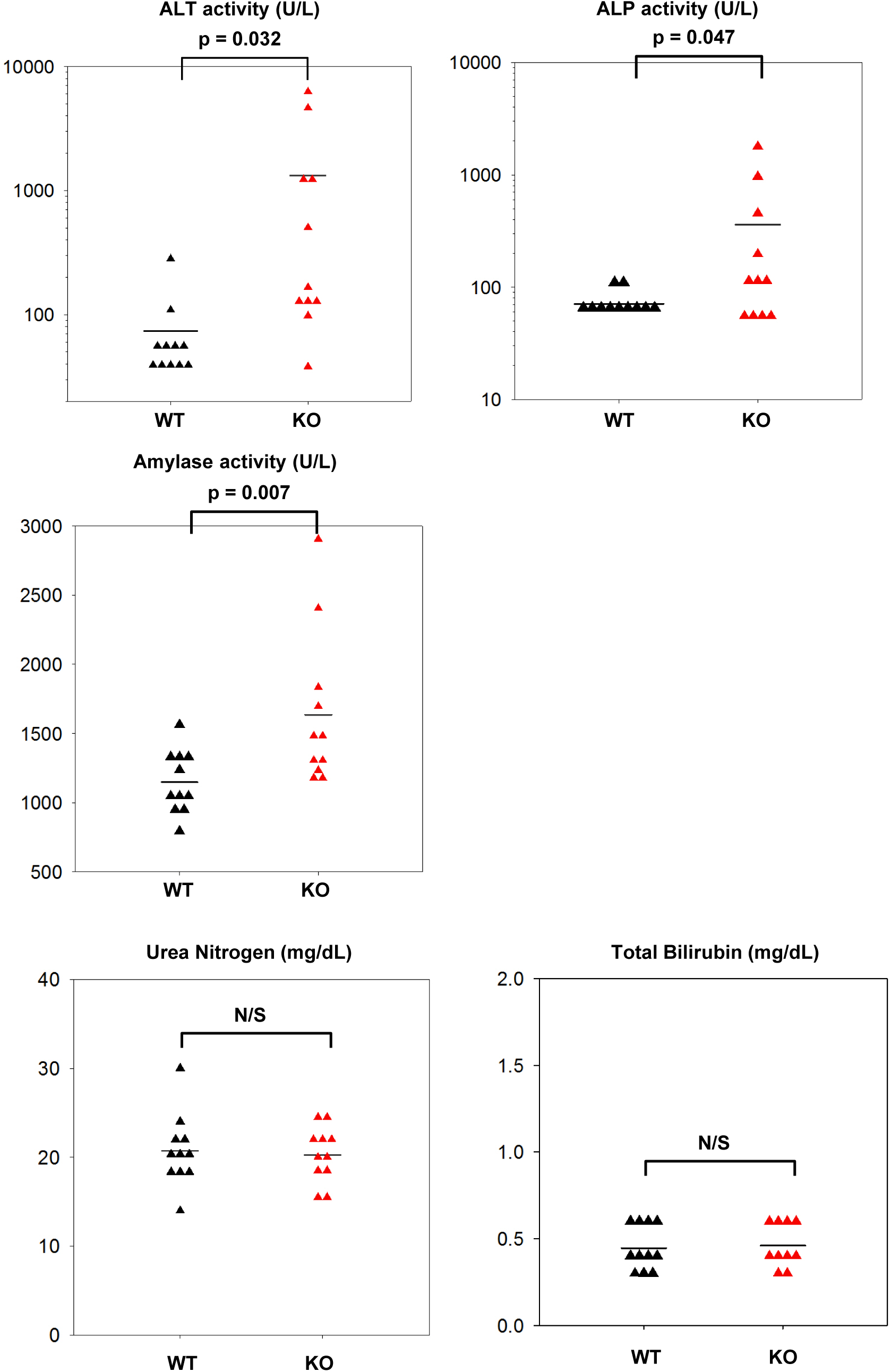
Srsf3 KO affects mouse liver function changes in the 1-year-old animals. Blood samples were collected from DENinjected Srsf3 WT and liver Srsf3 KO mice at a year of age and serum ALT, alkaline phosphatase, amylase activity, urea nitrogen and bilirubin were measured in a blood chemistry laboratory. A p-value was calculated by Student’s t-test. N/S, not significant.

**Fig. S5.**
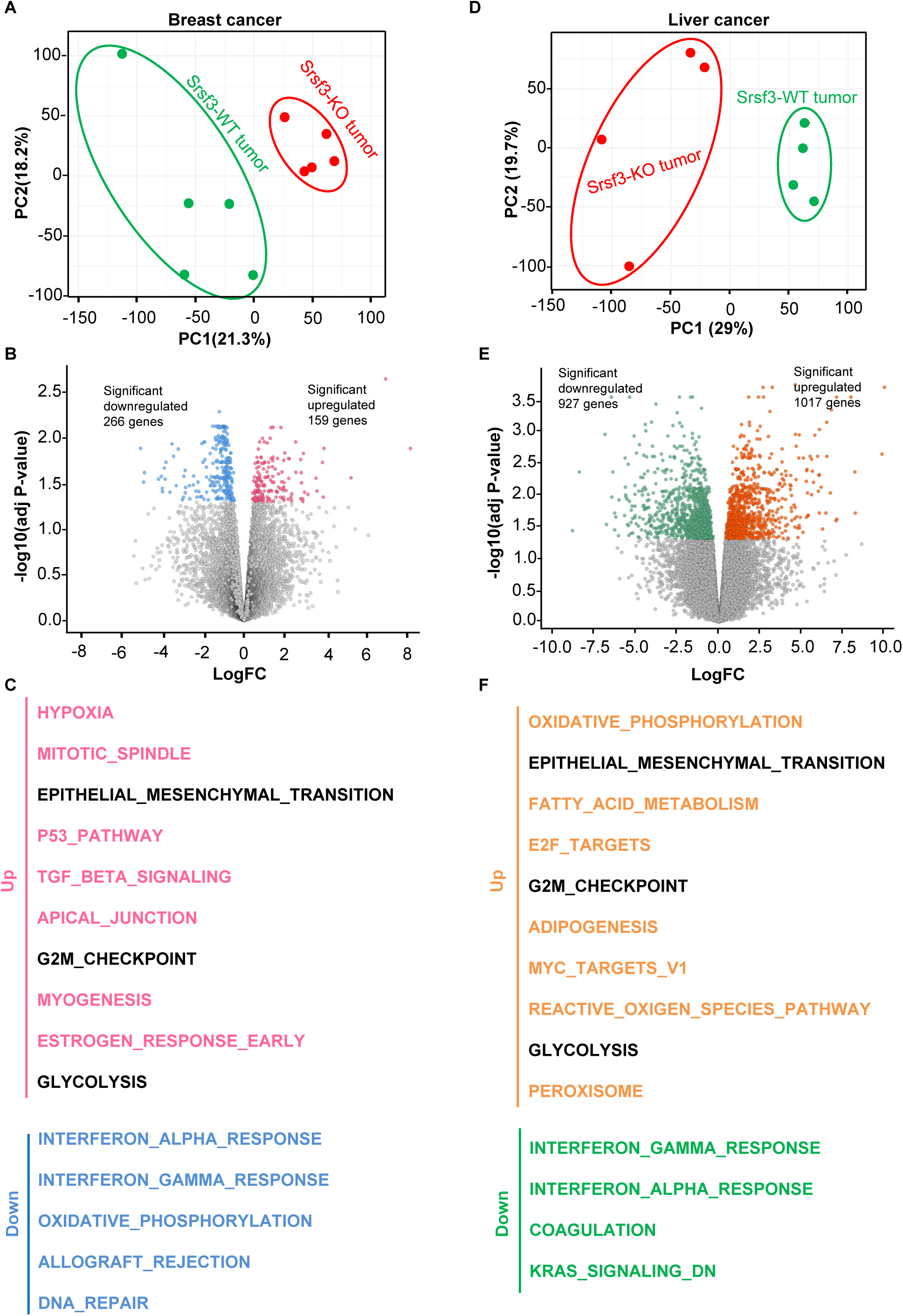
RNA-seq analysis of Srsf3 associated genes in Srsf3-WT and Srsf3-KO breast and liver cancers. Principal component analysis (PCA) of Srsf3 WT (green) and Srsf3 KO (red) breast cancer (A) and liver cancer (D) by RNA-seq analysis. Five mammary gland cancer tissues and four liver tcancer tissues were submitted to RNA-seq analysis in each group. (B and E) Volcano plot showing the down-regulated and upregulated genes (adjusted p ≤ 0.05) in Srsf3 KO cancer compared to Srsf3 WT cancer. (C and F) Top activated and inhibited pathways by Gene Set Enrichment Analysis (GSEA) performed with the Hallmark gene sets.

**Fig. S6.**
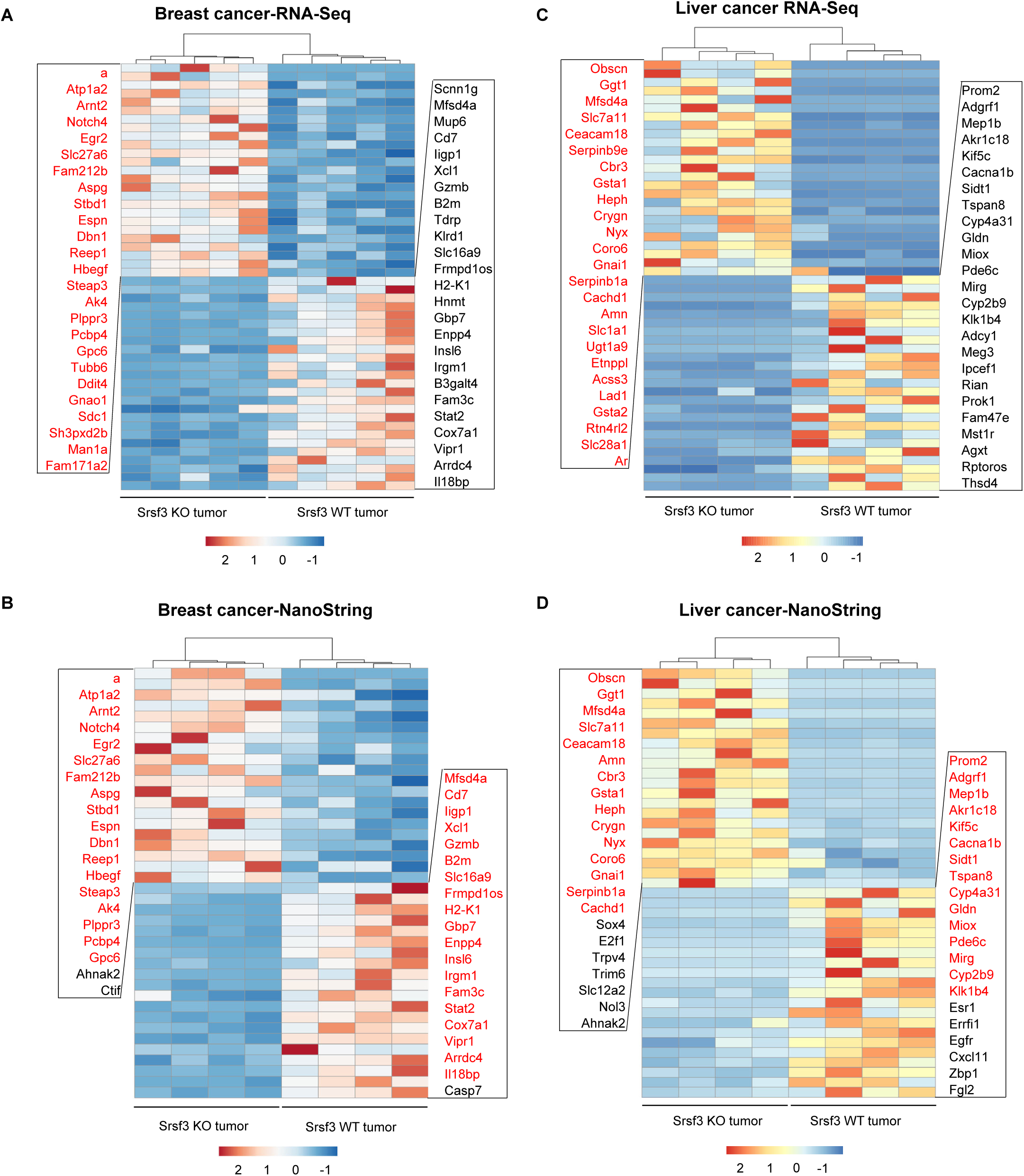
Identification of the top Srsf3 regulated gens by RNA-seq and validation by NanoString analysis in breast cancer and liver cancer. **(A** and **C)** Hierarchical clustering and heatmap of top 25 genes significantly upregulated and downregulated by Srsf3 KO in breast cancer (A) (Adjusted p ≤ 0.05, minRPKM >=2) and in liver cancer (C) (Adjusted p ≤ 0.05, minRPKM >=5) by RNA-seq analysis. (**B** and **D**) Selective validation by NanoString technology of the RNA-seq identified top genes up- or down-regulated by Srsf3 KO in breast cancer (**B**) and in liver cancer (**D**). Genes highlighted in red in heatmap are the genes identified by RNA-seq and validated by NanoString.

**Fig. S7.**
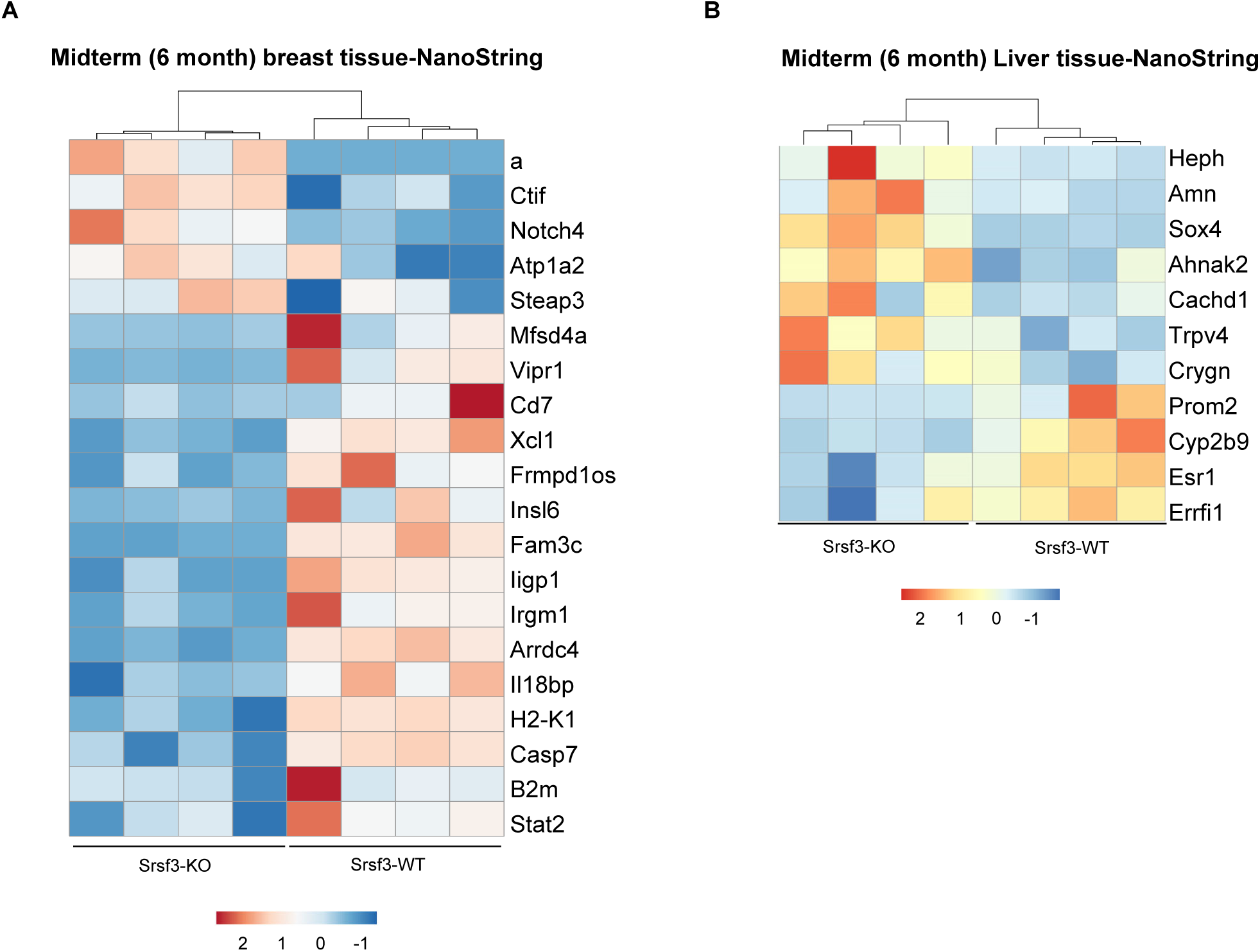
Srsf3 KO in breast and liver tissues affects differential expression of a subset of genes. Total RNA isolated from the indicated mouse tissues with or without Srsf3 KO at six months of mouse age were used for NanoString RNA analysis. Mice donated liver tissues for this study were also treated by DEN injected at day 15 of age. Heatmap shows differential expression of Srsf3 target genes identified in breast (A) and liver (B) tissues at this stage of animals.

**Fig. S8.**
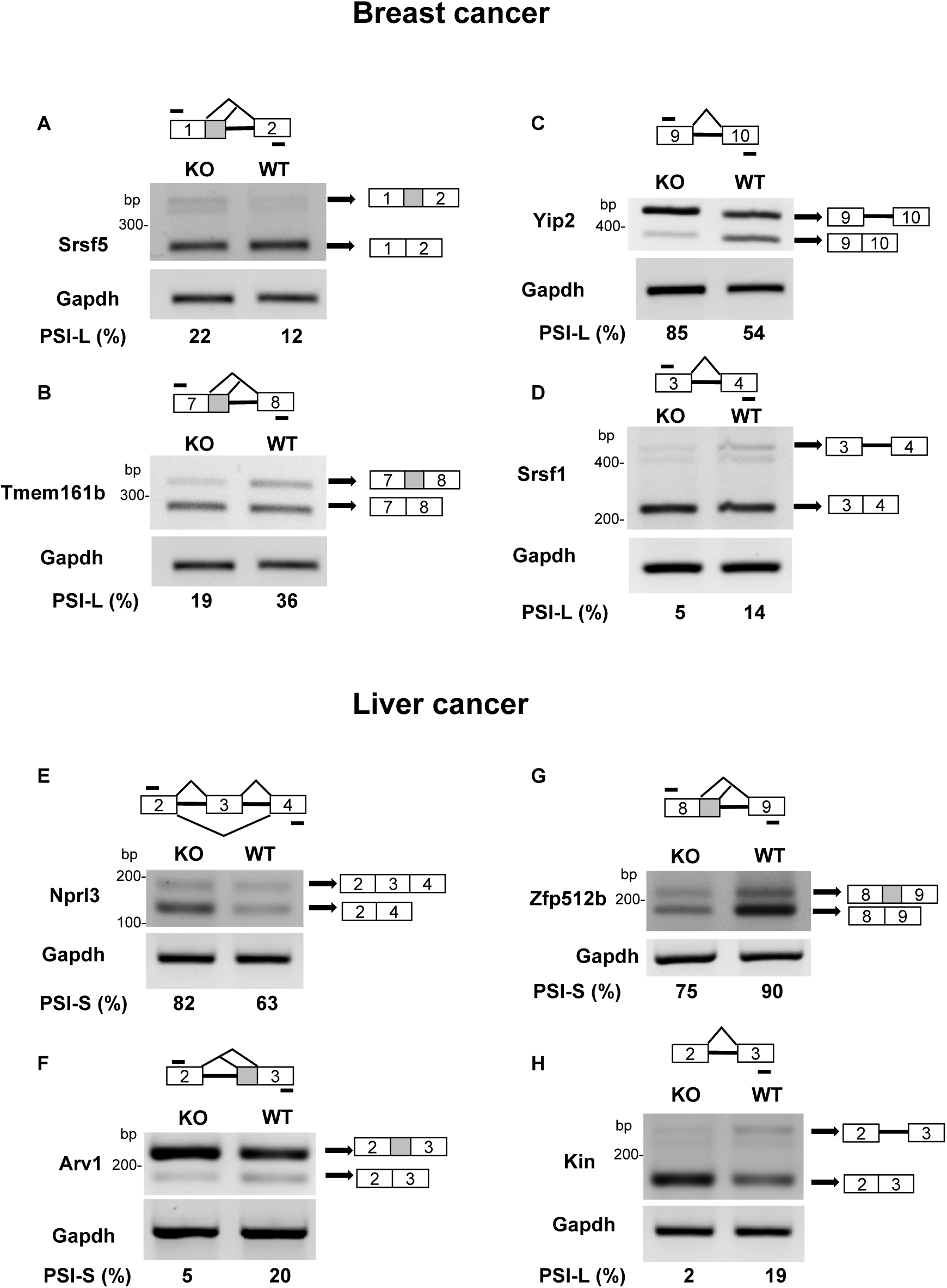
Validation of significant splicing events in differential expression of the Srsf3 target genes from Srsf3-KO to Srsf3-WT breast cancer (A-D) and liver cancer (E-H). Total RNA from the Srsf3-WT or srsf3-KO cancer tissues were analyzed by RT-PCR to validate the altered splicing events identified by by rMATS (p<=0.05, FDR<=0.05). (A-D) Srsf3 KO in breast cancer tissues alters alternative 5′ ss usage in Srsf5 exon 1 (A) and in Tmem161b exon 7 (B) and affects intron retention of Yip2 intron 9 (C) and Srsf1 intron 3 (D). (E-H) Srsf3 KO in liver cancer tissues decreases Nprl3 exon 3 skipping (E) but increases alternative 3′ ss usage in the Arv1exon 3 (F), alternative 5′ ss usage in the Zfp512b exon 8 (G), and Kin intron 2 retention (H). The primers used in RT-PCR are shown as bars above (forward primers) and below (reverse primers) each pre-mRNA diagram. GAPDH served as a loading control. PSI, percent spliced-in of the alternative exon(s) or splice site (% inclusion = inclusion/sum of inclusion + exclusion).

**Fig. S9.**
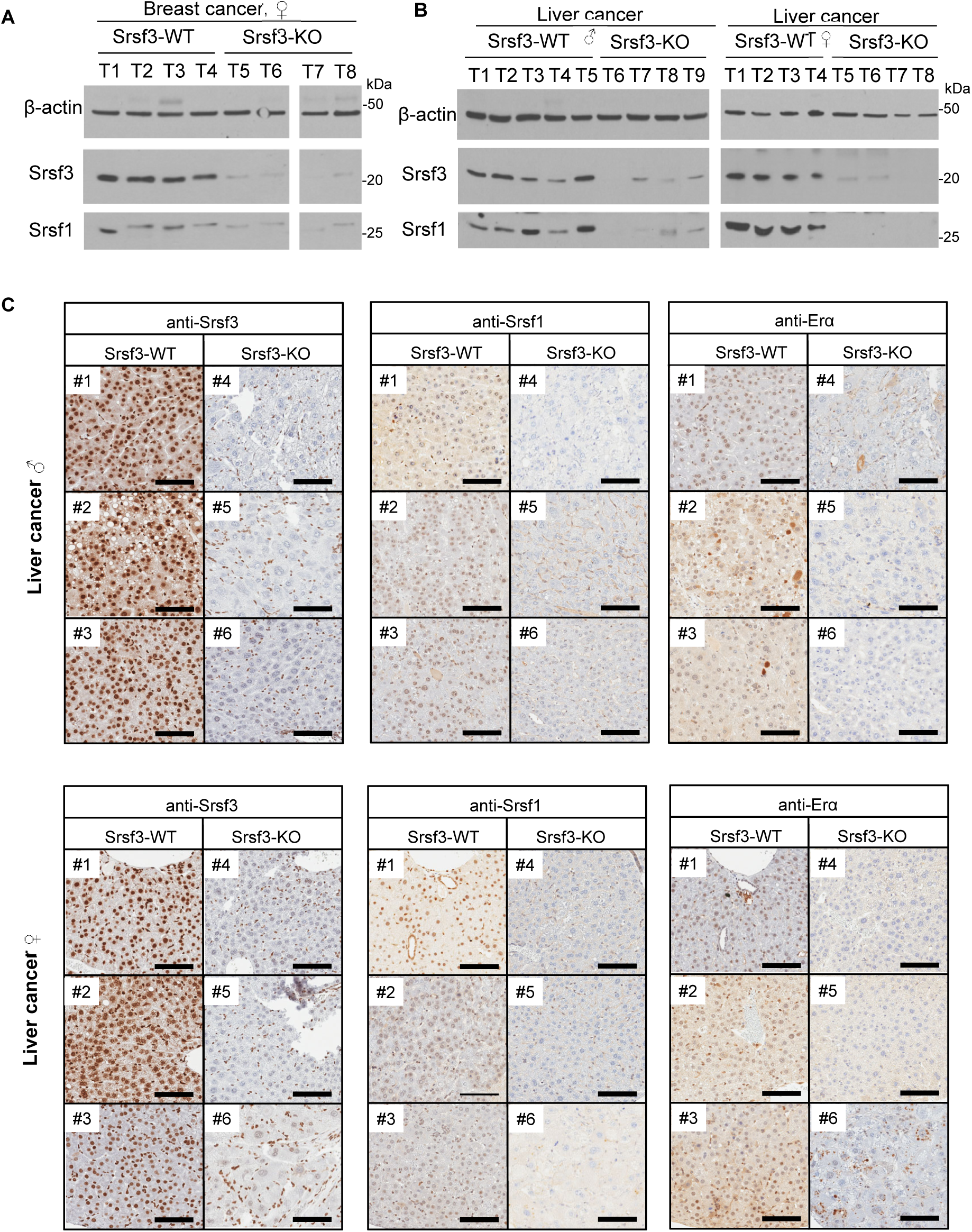
Srsf3 KO decreased SRSF1 protein expression in both breast and liver cancers. (A and B) knockout of Srsf3 expression in breast cancer (A) and liver cancer (B) led to reduce the protein expression of SRSF1 by Western blot analysis. Minimal four cancer tissues in each group were used for the assay. β-actin served as a loading control. (C) Immunohistology staining of Srsf3, Srsf1and Erα protein in both male and female SRSF3-WT and -KO liver cancer tissues. Three pairs of representative samples in both genders were shown. Scale bar, 100 μm.

**Fig. S10.**
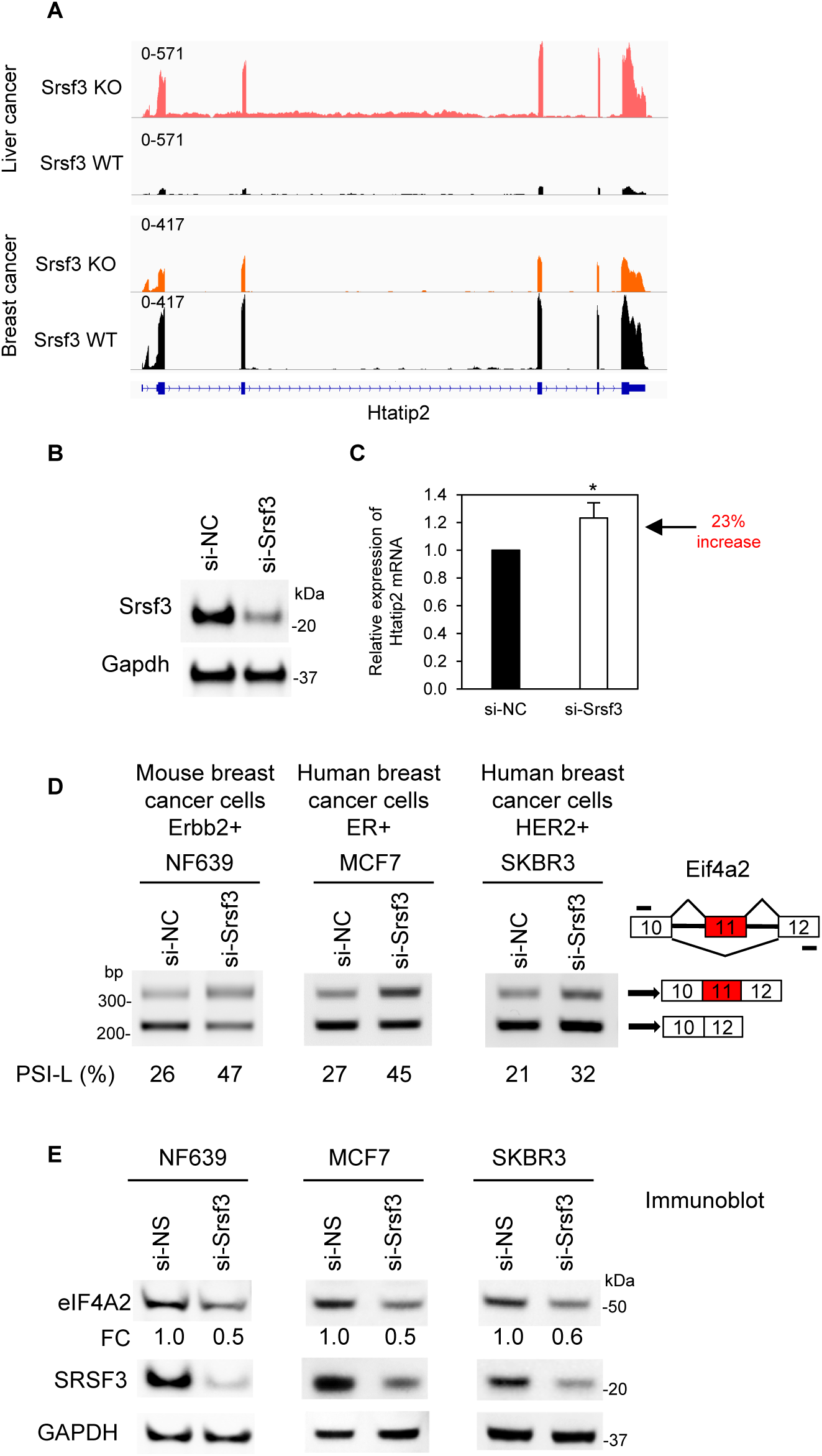
Srsf3 knockout promotes Htatip2 expression in liver cancer but reduces Htatip2 expression in breast cancer visualized by IGV (A). Srsf3 KD in Hepa 1-6 cells, a murine hepatoma cell line, led to increased expression of Htatip2. Hepa1-6 cells were transfected with 40 nM of Srsf3-specific siRNA (si-Srsf3) or non-targeting siRNA (si-NC) and harvested at 48h upon transfection for analysis. Srsf3 KD efficiency was confirmed by Western blot, with Gapdh serving as a sample loading control (B). (C) Srsf3 KD in Hepa 1-6 cells led to increased expression of Htatip2 examined by TaqMan RT-qPCR in three independent experiments. *, p< 0.05 by Student’s t-test.. (D-E) Knockdown (KD) of Srsf3/SRSF3 promotes exon 11 inclusion in Eif4a2 RNA splicing leading to reduction of eIF4A2 protein in mouse and human breast cancer cells. Total cell RNA was extracted from the cultured individual cell lines transfected with 40 nM of the indicated siRNAs and harvested at 48 h after transfection and used for RT-PCR assays with a forward primer from exon 10 and backward primer from exon 12 (D). Total cell extracts from corresponding cell lines prepared at the 48 h after siRNA transfection were immunoblotted for the protein expression of eIF4A2, SRSF3, and GAPDH with the corresponding antibodies (E).

**Fig. S11.**
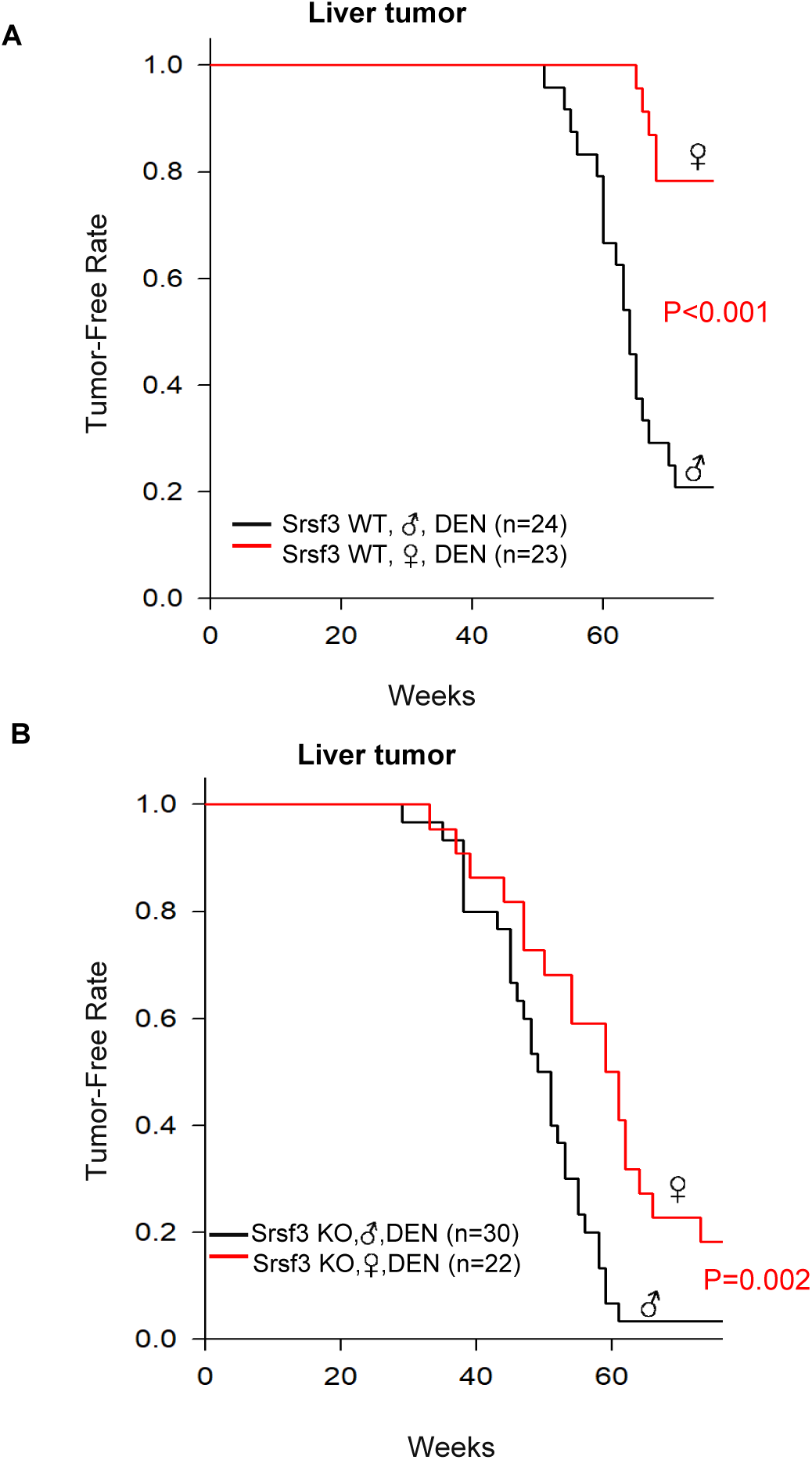
Gender disparity in DEN- and DEN+Srsf3-KO induced liver tumor formation between male and female mice. Kaplan-Meyer plots were applied for the incidence of palpable tumors in Srsf3-WT (A) and Srsf3-KO mice (B) after DEN injection on day 15 of age. N= number of animals in each group. P-values were determined by log-rank test between indicated groups.

**Fig. S12.**
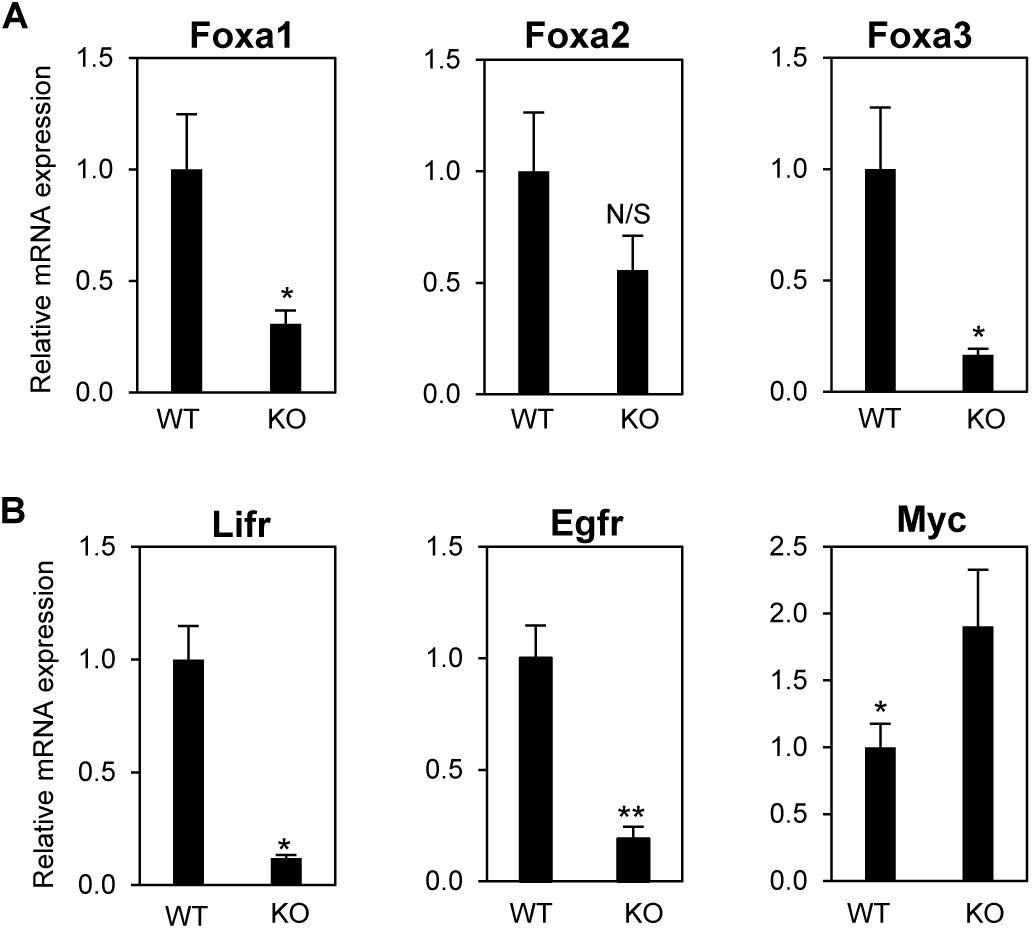
RNA expression of Foxa1, Foxa 2, Foxa 3 (A) and Lifr, Egfr, Myc (B) in Srsf3-WT and Srsf3 KO male liver tumors by quantitative RT-qPCR. Five samples were in each group. N/S, not significant, *, p < 0.05; **, p< 0.01 by Student’s t-test.

## Notes

### Competing Interest Statement

The authors have declared no competing interest.

## References

1. Bradley, R.K. and Anczuków, O. (2023) RNA splicing dysregulation and the hallmarks of cancer. Nat Rev Cancer, 23, 135–155.

2. Anczuków, O. and Krainer, A.R. (2016) Splicing-factor alterations in cancers. Rna, 22, 1285–1301.

3. David, C.J. and Manley, J.L. (2010) Alternative pre-mRNA splicing regulation in cancer: pathways and programs unhinged. Genes Dev, 24, 2343–2364.

4. Jia, R. and Zheng, Z.M. (2023) Oncogenic SRSF3 in health and diseases. Int J Biol Sci, 19, 3057–3076.

5. Jia, R., Ajiro, M., Yu, L., McCoy, P., Jr. and Zheng, Z.M. (2019) Oncogenic splicing factor SRSF3 regulates ILF3 alternative splicing to promote cancer cell proliferation and transformation. Rna, 25, 630–644.

6. Ajiro, M., Tang, S., Doorbar, J. and Zheng, Z.M. (2016) Serine/Arginine-Rich Splicing Factor 3 and Heterogeneous Nuclear Ribonucleoprotein A1 Regulate Alternative RNA Splicing and Gene Expression of Human Papillomavirus 18 through Two Functionally Distinguishable cis Elements. J Virol, 90, 9138–9152.

7. Ajiro, M., Jia, R., Yang, Y., Zhu, J. and Zheng, Z.M. (2016) A genome landscape of SRSF3-regulated splicing events and gene expression in human osteosarcoma U2OS cells. Nucleic Acids Res, 44, 1854–1870.

8. Wang, Z., Chatterjee, D., Jeon, H.Y., Akerman, M., Vander Heiden, M.G., Cantley, L.C. and Krainer, A.R. (2012) Exon-centric regulation of pyruvate kinase M alternative splicing via mutually exclusive exons. J Mol Cell Biol, 4, 79–87.

9. Jia, R., Liu, X., Tao, M., Kruhlak, M., Guo, M., Meyers, C., Baker, C.C. and Zheng, Z.M. (2009) Control of the papillomavirus early-to-late switch by differentially expressed SRp20. J Virol, 83, 167–180.

10. Majerciak, V., Lu, M., Li, X. and Zheng, Z.M. (2014) Attenuation of the suppressive activity of cellular splicing factor SRSF3 by Kaposi sarcoma-associated herpesvirus ORF57 protein is required for RNA splicing. Rna, 20, 1747–1758.

11. Lou, H., Neugebauer, K.M., Gagel, R.F. and Berget, S.M. (1998) Regulation of alternative polyadenylation by U1 snRNPs and SRp20. Mol Cell Biol, 18, 4977–4985.

12. Maciolek, N.L. and McNally, M.T. (2007) Serine/arginine-rich proteins contribute to negative regulator of splicing element-stimulated polyadenylation in rous sarcoma virus. J Virol, 81, 11208–11217.

13. Huang, Y. and Steitz, J.A. (2001) Splicing factors SRp20 and 9G8 promote the nucleocytoplasmic export of mRNA. Mol Cell, 7, 899–905.

14. Huang, Y., Gattoni, R., Stévenin, J. and Steitz, J.A. (2003) SR splicing factors serve as adapter proteins for TAP-dependent mRNA export. Mol Cell, 11, 837–843.

15. Auyeung, V.C., Ulitsky, I., McGeary, S.E. and Bartel, D.P. (2013) Beyond secondary structure: primary-sequence determinants license pri-miRNA hairpins for processing. Cell, 152, 844–858.

16. Kim, K., Nguyen, T.D., Li, S. and Nguyen, T.A. (2018) SRSF3 recruits DROSHA to the basal junction of primary microRNAs. Rna, 24, 892–898.

17. Fitzgerald, K.D. and Semler, B.L. (2011) Re-localization of cellular protein SRp20 during poliovirus infection: bridging a viral IRES to the host cell translation apparatus. PLoS Pathog, 7, e1002127.

18. Kim, J., Park, R.Y., Chen, J.K., Kim, J., Jeong, S. and Ohn, T. (2014) Splicing factor SRSF3 represses the translation of programmed cell death 4 mRNA by associating with the 5’-UTR region. Cell Death Differ, 21, 481–490.

19. Jumaa, H., Wei, G. and Nielsen, P.J. (1999) Blastocyst formation is blocked in mouse embryos lacking the splicing factor SRp20. Curr Biol, 9, 899–902.

20. Jia, R., Li, C., McCoy, J.P., Deng, C.X. and Zheng, Z.M. (2010) SRp20 is a proto-oncogene critical for cell proliferation and tumor induction and maintenance. Int J Biol Sci, 6, 806–826.

21. Tang, Y., Horikawa, I., Ajiro, M., Robles, A.I., Fujita, K., Mondal, A.M., Stauffer, J.K., Zheng, Z.M. and Harris, C.C. (2013) Downregulation of splicing factor SRSF3 induces p53β, an alternatively spliced isoform of p53 that promotes cellular senescence. Oncogene, 32, 2792–2798.

22. Ankö, M.L., Morales, L., Henry, I., Beyer, A. and Neugebauer, K.M. (2010) Global analysis reveals SRp20- and SRp75-specific mRNPs in cycling and neural cells. Nat Struct Mol Biol, 17, 962–970.

23. Kurokawa, K., Akaike, Y., Masuda, K., Kuwano, Y., Nishida, K., Yamagishi, N., Kajita, K., Tanahashi, T. and Rokutan, K. (2014) Downregulation of serine/arginine-rich splicing factor 3 induces G1 cell cycle arrest and apoptosis in colon cancer cells. Oncogene, 33, 1407–1417.

24. Jia, R., Zhang, S., Liu, M., Zhang, Y., Liu, Y., Fan, M. and Guo, J. (2016) HnRNP L is important for the expression of oncogene SRSF3 and oncogenic potential of oral squamous cell carcinoma cells. Sci Rep, 6, 35976.

25. Bray, F., Laversanne, M., Sung, H., Ferlay, J., Siegel, R.L., Soerjomataram, I. and Jemal, A. (2024) Global cancer statistics 2022: GLOBOCAN estimates of incidence and mortality worldwide for 36 cancers in 185 countries. CA Cancer J Clin, 74, 229–263.

26. Ke, H., Zhao, L., Zhang, H., Feng, X., Xu, H., Hao, J., Wang, S., Yang, Q., Zou, L., Su, X. et al. (2018) Loss of TDP43 inhibits progression of triple-negative breast cancer in coordination with SRSF3. Proc Natl Acad Sci U S A, 115, E3426–e3435.

27. Guo, L., Ke, H., Zhang, H., Zou, L., Yang, Q., Lu, X., Zhao, L. and Jiao, B. (2022) TDP43 promotes stemness of breast cancer stem cells through CD44 variant splicing isoforms. Cell Death Dis, 13, 428.

28. Sen, S., Jumaa, H. and Webster, N.J. (2013) Splicing factor SRSF3 is crucial for hepatocyte differentiation and metabolic function. Nat Commun, 4, 1336.

29. Kumar, D., Das, M., Sauceda, C., Ellies, L.G., Kuo, K., Parwal, P., Kaur, M., Jih, L., Bandyopadhyay, G.K., Burton, D. et al. (2019) Degradation of splicing factor SRSF3 contributes to progressive liver disease. J Clin Invest, 129, 4477–4491.

30. Sen, S., Langiewicz, M., Jumaa, H. and Webster, N.J. (2015) Deletion of serine/arginine-rich splicing factor 3 in hepatocytes predisposes to hepatocellular carcinoma in mice. Hepatology, 61, 171–183.

31. Yu, L., Majerciak, V., Jia, R. and Zheng, Z.M. (2023) Revisiting and corrections to the annotated SRSF3 (SRp20) gene structure and RefSeq sequences from the human and mouse genomes. Cell Insight, 2, 100089.

32. Muller, W.J., Sinn, E., Pattengale, P.K., Wallace, R. and Leder, P. (1988) Single-step induction of mammary adenocarcinoma in transgenic mice bearing the activated c-neu oncogene. Cell, 54, 105–115.

33. Naugler, W.E., Sakurai, T., Kim, S., Maeda, S., Kim, K., Elsharkawy, A.M. and Karin, M. (2007) Gender disparity in liver cancer due to sex differences in MyD88-dependent IL-6 production. Science, 317, 121–124.

34. Shen, S., Park, J.W., Huang, J., Dittmar, K.A., Lu, Z.X., Zhou, Q., Carstens, R.P. and Xing, Y. (2012) MATS: a Bayesian framework for flexible detection of differential alternative splicing from RNA-Seq data. Nucleic Acids Res, 40, e61.

35. Jumaa, H., Guénet, J.L. and Nielsen, P.J. (1997) Regulated expression and RNA processing of transcripts from the Srp20 splicing factor gene during the cell cycle. Mol Cell Biol, 17, 3116–3124.

36. Jumaa, H. and Nielsen, P.J. (1997) The splicing factor SRp20 modifies splicing of its own mRNA and ASF/SF2 antagonizes this regulation. Embo j, 16, 5077–5085.

37. Tan, X.Y., Li, Y.T., Li, H.H., Ma, L.X., Zeng, C.M., Zhang, T.T., Huang, T.X., Zhao, X.D. and Fu, L. (2023) WNT2-SOX4 positive feedback loop promotes chemoresistance and tumorigenesis by inducing stem-cell like properties in gastric cancer. Oncogene, 42, 3062–3074.

38. Liu, J., Qiu, J., Zhang, Z., Zhou, L., Li, Y., Ding, D., Zhang, Y., Zou, D., Wang, D., Zhou, Q. et al. (2021) SOX4 maintains the stemness of cancer cells via transcriptionally enhancing HDAC1 revealed by comparative proteomics study. Cell Biosci, 11, 23.

39. Hagiwara, M., Yasumizu, Y., Yamashita, N., Rajabi, H., Fushimi, A., Long, M.D., Li, W., Bhattacharya, A., Ahmad, R., Oya, M. et al. (2021) MUC1-C Activates the BAF (mSWI/SNF) Complex in Prostate Cancer Stem Cells. Cancer Res, 81, 1111–1122.

40. Gilchrist, C.L., Leddy, H.A., Kaye, L., Case, N.D., Rothenberg, K.E., Little, D., Liedtke, W., Hoffman, B.D. and Guilak, F. (2019) TRPV4-mediated calcium signaling in mesenchymal stem cells regulates aligned collagen matrix formation and vinculin tension. Proc Natl Acad Sci U S A, 116, 1992–1997.

41. Sato, T., Okumura, F., Ariga, T. and Hatakeyama, S. (2012) TRIM6 interacts with Myc and maintains the pluripotency of mouse embryonic stem cells. J Cell Sci, 125, 1544–1555.

42. Jaworska, A.M., Wlodarczyk, N.A., Mackiewicz, A. and Czerwinska, P. (2020) The role of TRIM family proteins in the regulation of cancer stem cell self-renewal. Stem Cells, 38, 165–173.

43. Karni, R., de Stanchina, E., Lowe, S.W., Sinha, R., Mu, D. and Krainer, A.R. (2007) The gene encoding the splicing factor SF2/ASF is a proto-oncogene. Nat Struct Mol Biol, 14, 185–193.

44. Anczuków, O., Rosenberg, A.Z., Akerman, M., Das, S., Zhan, L., Karni, R., Muthuswamy, S.K. and Krainer, A.R. (2012) The splicing factor SRSF1 regulates apoptosis and proliferation to promote mammary epithelial cell transformation. Nat Struct Mol Biol, 19, 220–228.

45. Anczuków, O., Akerman, M., Cléry, A., Wu, J., Shen, C., Shirole, N.H., Raimer, A., Sun, S., Jensen, M.A., Hua, Y. et al. (2015) SRSF1-Regulated Alternative Splicing in Breast Cancer. Mol Cell, 60, 105–117.

46. Du, J.X., Luo, Y.H., Zhang, S.J., Wang, B., Chen, C., Zhu, G.Q., Zhu, P., Cai, C.Z., Wan, J.L., Cai, J.L. et al. (2021) Splicing factor SRSF1 promotes breast cancer progression via oncogenic splice switching of PTPMT1. J Exp Clin Cancer Res, 40, 171.

47. Ito, M., Jiang, C., Krumm, K., Zhang, X., Pecha, J., Zhao, J., Guo, Y., Roeder, R.G. and Xiao, H. (2003) TIP30 deficiency increases susceptibility to tumorigenesis. Cancer Res, 63, 8763–8767.

48. Pecha, J., Ankrapp, D., Jiang, C., Tang, W., Hoshino, I., Bruck, K., Wagner, K.U. and Xiao, H. (2007) Deletion of Tip30 leads to rapid immortalization of murine mammary epithelial cells and ductal hyperplasia in the mammary gland. Oncogene, 26, 7423–7431.

49. Li, A., Zhang, C., Gao, S., Chen, F., Yang, C., Luo, R. and Xiao, H. (2013) TIP30 loss enhances cytoplasmic and nuclear EGFR signaling and promotes lung adenocarcinogenesis in mice. Oncogene, 32, 2273–2281, 2281e.2271-2212.

50. Kalra, M., Mayes, J., Assefa, S., Kaul, A.K. and Kaul, R. (2008) Role of sex steroid receptors in pathobiology of hepatocellular carcinoma. World J Gastroenterol, 14, 5945–5961.

51. Shimizu, I., Yasuda, M., Mizobuchi, Y., Ma, Y.R., Liu, F., Shiba, M., Horie, T. and Ito, S. (1998) Suppressive effect of oestradiol on chemical hepatocarcinogenesis in rats. Gut, 42, 112–119.

52. Nakatani, T., Roy, G., Fujimoto, N., Asahara, T. and Ito, A. (2001) Sex hormone dependency of diethylnitrosamine-induced liver tumors in mice and chemoprevention by leuprorelin. Jpn J Cancer Res, 92, 249–256.

53. Li, Z., Tuteja, G., Schug, J. and Kaestner, K.H. (2012) Foxa1 and Foxa2 are essential for sexual dimorphism in liver cancer. Cell, 148, 72–83.

54. Shachaf, C.M., Kopelman, A.M., Arvanitis, C., Karlsson, A., Beer, S., Mandl, S., Bachmann, M.H., Borowsky, A.D., Ruebner, B., Cardiff, R.D. et al. (2004) MYC inactivation uncovers pluripotent differentiation and tumour dormancy in hepatocellular cancer. Nature, 431, 1112–1117.

55. Derynck, R., Akhurst, R.J. and Balmain, A. (2001) TGF-beta signaling in tumor suppression and cancer progression. Nat Genet, 29, 117–129.

56. Yang, L. and Moses, H.L. (2008) Transforming growth factor beta: tumor suppressor or promoter? Are host immune cells the answer? Cancer Res, 68, 9107–9111.

57. Perkins, N.D. (2012) The diverse and complex roles of NF-κB subunits in cancer. Nat Rev Cancer, 12, 121–132.

58. Tesio, M., Trinquand, A., Macintyre, E. and Asnafi, V. (2016) Oncogenic PTEN functions and models in T-cell malignancies. Oncogene, 35, 3887–3896.

59. Shojaee, S., Chan, L.N., Buchner, M., Cazzaniga, V., Cosgun, K.N., Geng, H., Qiu, Y.H., von Minden, M.D., Ernst, T., Hochhaus, A., et al. (2016) PTEN opposes negative selection and enables oncogenic transformation of pre-B cells. Nat Med, 22, 379–387.

60. Wimmer, P., Berscheminski, J., Blanchette, P., Groitl, P., Branton, P.E., Hay, R.T., Dobner, T. and Schreiner, S. (2016) PML isoforms IV and V contribute to adenovirus-mediated oncogenic transformation by functionally inhibiting the tumor-suppressor p53. Oncogene, 35, 69–82.

61. Renner, F., Moreno, R. and Schmitz, M.L. (2010) SUMOylation-dependent localization of IKKepsilon in PML nuclear bodies is essential for protection against DNA-damage-triggered cell death. Mol Cell, 37, 503–515.

62. Wu, S.Y., Lee, C.F., Lai, H.T., Yu, C.T., Lee, J.E., Zuo, H., Tsai, S.Y., Tsai, M.J., Ge, K., Wan, Y. et al. (2020) Opposing Functions of BRD4 Isoforms in Breast Cancer. Mol Cell, 78, 1114–1132.e1110.

63. Shen, S., Wang, Y., Wang, C., Wu, Y.N. and Xing, Y. (2016) SURVIV for survival analysis of mRNA isoform variation. Nat Commun, 7, 11548.

64. Kumar, D., Das, M., Oberg, A., Sahoo, D., Wu, P., Sauceda, C., Jih, L., Ellies, L.G., Langiewicz, M.T., Sen, S. et al. (2022) Hepatocyte Deletion of IGF2 Prevents DNA Damage and Tumor Formation in Hepatocellular Carcinoma. Adv Sci (Weinh), 9, e2105120.

65. Wang, H., Lekbaby, B., Fares, N., Augustin, J., Attout, T., Schnuriger, A., Cassard, A.M., Panasyuk, G., Perlemuter, G., Bieche, I. et al. (2019) Alteration of splicing factors’ expression during liver disease progression: impact on hepatocellular carcinoma outcome. Hepatol Int, 13, 454–467.

66. Sherr, C.J. and Roberts, J.M. (1995) Inhibitors of mammalian G1 cyclin-dependent kinases. Genes Dev, 9, 1149–1163.

67. Liggett, W.H., Jr. and Sidransky, D. (1998) Role of the p16 tumor suppressor gene in cancer. J Clin Oncol, 16, 1197–1206.

68. Chung, C.H., Zhang, Q., Kong, C.S., Harris, J., Fertig, E.J., Harari, P.M., Wang, D., Redmond, K.P., Shenouda, G., Trotti, A. et al. (2014) p16 protein expression and human papillomavirus status as prognostic biomarkers of nonoropharyngeal head and neck squamous cell carcinoma. J Clin Oncol, 32, 3930–3938.

69. Romagosa, C., Simonetti, S., López-Vicente, L., Mazo, A., Lleonart, M.E., Castellvi, J. and Ramon y Cajal, S. (2011) p16(Ink4a) overexpression in cancer: a tumor suppressor gene associated with senescence and high-grade tumors. Oncogene, 30, 2087–2097.

70. Wang, X., Meyers, C., Guo, M. and Zheng, Z.M. (2011) Upregulation of p18Ink4c expression by oncogenic HPV E6 via p53-miR-34a pathway. Int J Cancer, 129, 1362–1372.

71. Solomon, D.A., Kim, J.S., Jenkins, S., Ressom, H., Huang, M., Coppa, N., Mabanta, L., Bigner, D., Yan, H., Jean, W. et al. (2008) Identification of p18 INK4c as a tumor suppressor gene in glioblastoma multiforme. Cancer Res, 68, 2564–2569.

72. Abbas, T. and Dutta, A. (2009) p21 in cancer: intricate networks and multiple activities. Nat Rev Cancer, 9, 400–414.

73. Yang, H., Qin, G., Luo, Z., Kong, X., Gan, C., Zhang, R. and Jiang, W. (2022) MFSD4A inhibits the malignant progression of nasopharyngeal carcinoma by targeting EPHA2. Cell Death Dis, 13, 332.

74. Bhat, M., Robichaud, N., Hulea, L., Sonenberg, N., Pelletier, J. and Topisirovic, I. (2015) Targeting the translation machinery in cancer. Nat Rev Drug Discov, 14, 261–278.

